# Non-invasive imaging of cell-based therapies using acoustic reporter genes

**DOI:** 10.1101/2024.11.01.621111

**Authors:** Shirin Shivaei, Ann Liu, Mohamad H. Abedi, Julio Revilla, George Daghlian, Isabella U. Hurvitz, Hao Shen, Margaret B. Swift, Mikhail G. Shapiro

**Affiliations:** Division of Biology and Biological Engineering, California Institute of Technology; Pasadena, CA 91125, USA; Division of Chemistry and Chemical Engineering, California Institute of Technology; Pasadena, CA 91125, USA; Andrew and Peggy Cherng Department of Medical Engineering, California Institute of Technology; Pasadena, CA 91125, USA; Howard Hughes Medical Institute; Pasadena, CA 91125, USA

## Abstract

Cell-based therapies are a major emerging category of medicine. The ability of engineered cells to traffic to and function at specific anatomical locations is a major aspect of their performance. However, there is a lack of non-invasive, non-ionizing, cost-accessible methods to track these therapies inside the body and ensure proper function. Here, we establish a platform for *in vivo* imaging of primary cell therapies using ultrasound – a ubiquitously accessible technology for high-resolution non-invasive imaging. We introduce and optimize a lentiviral delivery system to express acoustic reporter genes based on gas vesicles in mammalian cells such as T cells, showing that this results in robust ultrasound contrast. Additionally, we develop genetic circuits making it possible to monitor T cell activation via activity-dependent promoters. We apply this technology to primary human T cells, using it to non-invasively track their accumulation and proliferation as a targeted therapy in a mouse tumor xenograft model and compare it to invasive, terminal measures such as immunohistology. By making it possible to visualize cell-based therapies and their function inside opaque living organs with unprecedented resolution and accessibility, this technology has the potential to significantly accelerate their development and effective use.

## INTRODUCTION

Cell-based therapies are making breakthrough progress in treating complex diseases such as cancer and autoimmune disease. However, predicting and monitoring their *in vivo* dynamics, including tissue infiltration, proliferation and off-target activity, remain challenging and necessitate the development of more advanced *in vivo* imaging methods. Existing imaging techniques used in research, such as luminescent reporters^1^, provide poor spatial resolution due to light scattering, while clinical probes based on positron emission tomography require specialized radioligand synthesis and imaging equipment, including just-in-time synthesis and delivery of radiotracers, and involve exposure to ionizing radiation^2–6^. Improved imaging techniques are essential for understanding the mechanisms of cell-based therapies, optimizing treatment protocols, and ensuring precise targeting of therapeutic cells to improve their efficacy and reduce off-target effects, ultimately accelerating the development of new cell-based treatments.

Ultrasound presents a promising modality to track cell therapies, as it is affordable, widely available in medical settings, and capable of large-scale scanning of deep organs with spatial resolution on the order of 100 micrometers^7^. To enable *in vivo* imaging of cell therapies with ultrasound, one would ideally use reporter genes rather than synthetic labels, as this would allow the tracking of proliferating cells without signal dilution and the sensing of specific cellular states via promoter-driven expression.

Here we introduce a method to track cell therapies with ultrasound using acoustic reporter genes based on gas vesicles (GVs). GVs are microbially derived air-filled protein nanostructures that scatter sound waves, enabling their use as reporter genes for ultrasound imaging in mammalian cells^8–10^ and probiotics^11,12,9,13^. However, previous mammalian uses of GVs have been limited to chemical transfections of the GV gene cluster, restricting their use to transfection-compatible cell lines. In fact, to date there has been no demonstration of GV gene expression in primary cells due to the difficulty of delivering the eight or more genes encoding GVs, at appropriate stoichiometries, to such cells using compatible methods such as viral delivery.

To overcome this challenge, we designed, optimized and extensively characterized a lentiviral vector architecture capable of delivering the complete set of GV-encoding genes into mammalian cells, resulting in robust ultrasound contrast. We showed that GV expression can be driven by a chemically inducible promoter or connected to a gene circuit engineered to respond to cell activity, permitting the imaging of receptor-driven immune cell activation. We used our lentiviral expression system to produce GV-expressing primary human T cells and used ultrasound to monitor their accumulation *in vivo* as an adoptively transferred cell therapy in mouse tumor xenografts, as corroborated by immunohistochemistry. We further show that intratumoral ultrasound signal from GV-expressing CAR-T cells provides an early, quantitative indicator of therapeutic efficacy, preceding optical or volumetric measurements of tumor response. These results position virally-encoded GVs as a prime option for ultrasound imaging of cell-based therapies.

## RESULTS

### Optimized lentiviral vector design enhances GV gene cluster delivery and expression in mammalian cells

To enable their use as acoustic reporters in primary cells, we sought to package the genes encoding GVs into lentiviral vectors – the most established delivery vehicle in clinical cell engineering^14^. Lentiviral packaging and transduction efficiency depend heavily on transgene size, with a packaging limit of 8-10 kilobases (kb) and reduced titers with larger payloads in this range^15^. The smallest GV gene cluster expressed in mammalian cells, derived from *Anabaena flos-aquae*, comprises 8 genes, which take up 4.5 kb without promoters and other genetic elements required for polycistronic expression^9^. To establish a strategy for efficient lentiviral GV expression, we decided to compare three vector architectures in which the genes are combined across one, two or three co-transduced viral constructs. We then transduced HEK293T cells with each viral combination at different multiplicities of infection (MOIs) and measured the ultrasound contrast of the cells embedded in hydrogel phantoms (**Fig. 1a**). We used two nonlinear ultrasound imaging methods optimized for GV detection – BURST^16^ and xAM^17^. The former makes use of the strong nonlinear scattering produced immediately after collapsing GVs with a high-pressure pulse, while the latter non-destructively captures nonlinear scattering arising from reversible buckling of the GVs with each cycle of the transmitted sound wave.

**Figure 1:**
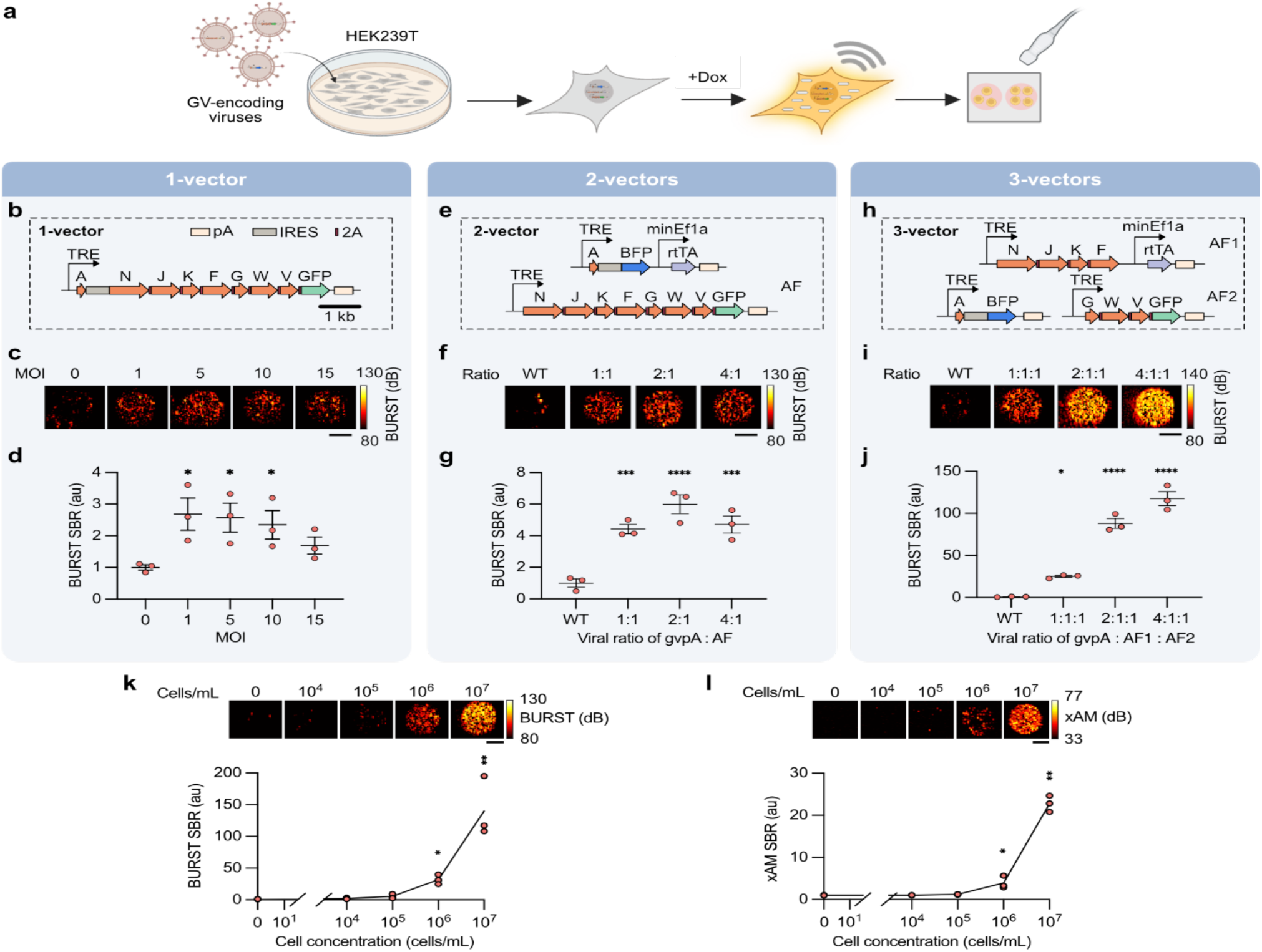
Design and optimization of multi-vector lentiviral delivery systems for robust GV expression and ultrasound imaging. **a**, Schematic of the viral production and screening process. HEK293T cells are transduced with lentiviral vectors encoding GV genes, followed by doxycycline-induced GV expression for 72 hours. Cells are detached and embedded in agarose phantoms for ultrasound imaging. **b**, Schematic of a single lentiviral vector encoding all 8 GV genes. **c**, Representative BURST ultrasound images of HEK-TetON cells transduced with the single-vector virus at various multiplicities of infection (MOIs). **d**, Quantification of BURST signal-to-background ratio (SBR) in cells transduced with the single-vector virus, normalized to non-transduced cells. **e**, Schematic of two lentiviral vectors encoding GV genes and the rtTA transactivator. **f**, Representative BURST images of HEK293T cells transduced with the two-vector system at a total MOI of 10, with varying ratios of *gvpA* virus to assembly factor (AF) virus. **g**, Quantification of BURST SBR in cells from transduced with (**e**). **h**, Schematic of a three-vector system encoding the GV genes. **i**, Representative BURST images of HEK293T cells transduced with the three-vector system at a total MOI of 10, with varying ratios of *gvpA* to the first assembly factor (AF1) virus and the second assembly factor (AF2) virus. **j**, Quantification of BURST SBR in cells transduced with (**h**). **k**, Representative BURST images (top) and SBR (bottom) of HEK293T cells transduced with the three-vector system at a 4:1:1 ratio and sorted by fluorescence, shown as a function of cell concentration. **l**, Representative xAM images (top) and corresponding SBR (bottom) of the same cells as panel (**k**), shown as a function of cell concentration. For panels (**d, g, j**), statistical comparisons to non-transduced controls were performed using Fisher’s least significant difference (LSD) test. For panels (**k-l**), comparisons were made against the “no cells” control using Kruskal-Wallis test. P-values from left to right: (**d**): 0.0017, 0.0165, 0.0333; (**g**): 0.0006, <0.0001, 0.0004; (**j**): 0.0107, <0.0001, <0.0001; (**k**): 0.0137, 0.001; (**l**): 0.0285, 0.0026. T-values from left to right: (**d**): 3.079, 2.877, 2.467; (**g**): 5.405, 7.878, 5.859; (**j**): 3.308, 11.98, 16.04; Df (**d**): 10; (**g**): 8; (**j**): 8; Z-values from left to right: (**k**): 2.465, 3.286; (**l**): 2.191, 3.012. Significance levels: \**p* < 0.05, \*\**p* < 0.01, \*\*\**p* < 0.001, \*\*\*\**p* < 0.0001; nonsignificant data points are not marked. For (**d, g, j**), error bars represent mean ± s.e.m. of N = 3 biological replicates. Each data point reflects the arithmetic mean of N = 2 technical replicates, unless noted otherwise. For (**k, l**), N = 3 replicates from independent dilution series. Scale bars for all ultrasound images represent 1 mm. au: arbitrary units. Dox: doxycycline.

In our first design, we used a single lentiviral backbone to encode the main structural protein *gvpA* and all seven GV assembly factor genes (*gvpN-V*), along with a fluorescent marker (GFP), under a TRE promoter (**Fig. 1b**). This promoter, used in conjunction with an rtTA transactivator, drives expression in the presence of doxycycline (Dox), allowing GV production to be timed with imaging to minimize metabolic burden. We linked *gvpA* to the assembly factors (AFs) via an internal ribosome entry site (IRES), while the AF genes were themselves polycistronically linked using self-cleaving 2A peptides (iterating through P2A, T2A, E2A, and F2A, to minimize the risk of homology-induced genetic recombination). The total packaging size was 8.8 kb. We transduced a HEK-TetON cell line with this virus, while varying the MOI from 1 to 15. We observed BURST ultrasound contrast at all MOIs, with the strongest signal at MOIs of 1, 5, and 10 and decreasing at MOI of 15 (**Fig. 1c-d**). The xAM contrast was strongest at MOI of 5 and decreased at MOIs of 10 and 15 (**Extended Data Fig. 1a**).

The highest MOIs also showed reduced infectivity via flow cytometry, suggesting toxicity from high viral burden (**Extended Data Fig. 2a-b**). These results show that it is possible to express GVs in mammalian cells using viral vectors, and that the entire GV gene cluster can fit into a single lentiviral construct. While recognizing these important milestones, we hypothesized that breaking the construct up into two or three smaller inserts would allow more efficient viral packaging and transduction, provide flexibility to control gene stoichiometry, and make room to include the rtTA transactivator to enable expression in previously unengineered cells.

Thus, our second design comprised two vectors: one encoding the main structural protein *gvpA* and rtTA, and the second encoding all the AFs, *gvpN-V* (**Fig. 1e**). These vector inserts had lengths of 5.1 and 8 kb, respectively. This two-vector design enabled us to adjust the copy number of *gvpA* relative to the AFs, a variable shown to impact GV expression in mammalian cells^9^. Transducing a HEK293T cell line, we maintained a total MOI of 10 while varying the MOI ratio between the *gvpA* and the AF vectors. We found that all ratios produced significant ultrasound contrast in BURST (**Fig. 1f-g**) and xAM (**Extended Data Fig. 1b**), peaking at 2:1 and 4:1 ratios, respectively. Transduction was similar across all ratios, remaining above 40%, and the expression of BFP and GFP followed the ratios of the *gvpA* and AF vectors as expected (**Extended Data Fig. 2c-d**).

In our third design, we split the seven AF genes across two vectors, grouping *gvpNJKF* (AF1) and *gvpGWV* (AF2), respectively, with packaging sizes of 6.7 and 4.8 kb. We encoded *gvpA* on its own third virus with an insert length of 4 kb (**Fig. 1h**). The rtTA was included on the AF1 construct. Using a constant total MOI of 10 while testing three ratios of the *gvpA* vector relative to the two AF vectors, we observed much stronger BURST contrast compared to the one- and two-vector designs (**Fig. 1i-j**), with the highest signal-to-background ratio (SBR) at the 4:1:1 ratio, measured to be 118 ± 8.4 (mean ± s.e.m.). The xAM signal was similarly improved and increased with more *gvpA* (**Extended Data Fig. 1c**). Transduction efficiency was comparable to the 2-vector design (**Extended Data Fig. 2e-f**). Cell viability was not affected by viral transduction or the induction of GV expression (**Extended Data Fig. 2g**). These results demonstrate that a split-vector approach comprising smaller inserts and more flexibility in ratiometric expression provides the best performance for expressing the polycistronic GV complex.

Using our best-performing three-vector transduction system (4:1:1), we quantified ultrasound contrast as a function of cell concentration after sorting for triple-transduced cells. Significant signal from the cells could be detected at a concentration of 1e6 cells per milliliter with BURST (**Fig. 1k**) and with xAM (**Fig. 1l**). This concentration corresponds to approximately 4 cells per image voxel (where each voxel is 0.004 mm^3^) and a volume fraction of approximately 0.9%. Thus, we concluded that viral vector delivery enables highly sensitive imaging of GV-expressing cells and adopted the three-vector design for the rest of our experiments.

### Lentiviral GV delivery system is adaptable for robust transduction and ultrasound imaging of immune cells

We next asked if our lentiviral GV delivery architecture could be adapted to image immune cells, starting with the Jurkat T cell line, which is commonly used to model lymphocyte-based cell therapies (**Fig. 2a**). Since Jurkats are more difficult to transduce than HEK239T cells^18^, we hypothesized that they would require higher viral doses to achieve similar levels of transduction. We therefore incubated the Jurkat cells with total MOIs of 30 and 60 (as defined based on functional tittering in HEK cells), varying the *gvpA*-to-AF1 and AF2 ratio within each MOI. While both MOIs resulted in appreciable ultrasound signal at certain ratios, the most consistent BURST contrast was produced at MOI of 60, with the 4:1:1 ratio showing the highest BURST SBR at 5.94 ± 1.64 (**Fig. 2b-c**), and the strongest signal in xAM (**Extended Data Fig. 3a-b**). As expected, cells transduced with the higher MOI had a larger number of triple-transduced cells (**Extended Data Fig. 4a**) and higher levels of GFP and BFP expression (**Extended Data Fig. 4b-c**).

**Figure 2:**
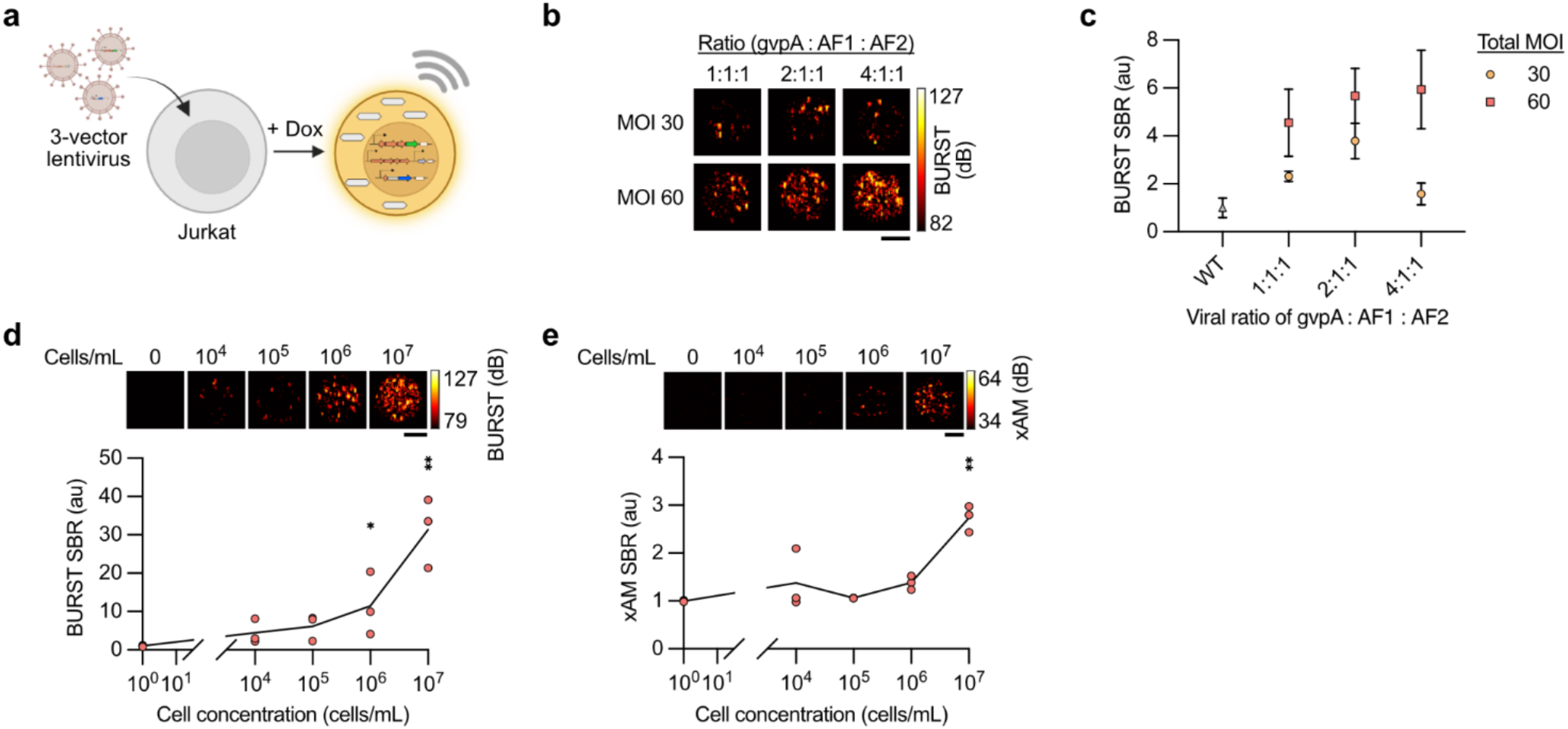
Adaptability of lentiviral GV gene delivery and ultrasound imaging of immune cells. **a**, Schematic of doxycycline-inducible GV expression in Jurkat cells transduced with three lentiviral vectors encoding the GV gene cluster. **b**, Representative BURST ultrasound images of Jurkat cells transduced with different total MOIs and three ratios of *gvpA* to assembly factor (AF1 and AF2) viruses. **c**, Quantified BURST signal in Jurkat cells from panel (**b**), normalized to wild-type (WT) Jurkat cells. **d**, Representative BURST images (top) and corresponding signal quantification (bottom) of Jurkat cells transduced with the three-vector lentivirus system (MOI 60) at a 4:1:1 ratio. Cells were sorted by fluorescence expression and imaged at varying cell concentrations to measure imaging detection limit. **e**, Representative xAM images (top) and signal quantification (bottom) of the same sorted cell line from panel (**d**), shown as a function of cell concentration. Statistical comparisons for panel (**c**) were performed using Fisher’s LSD test comparing each condition to WT. Statistics for MOI 60 from left to right: p = 0.0241, 0.005, 0.0034; t = 2.529, 3.328, 3.519; df = 14 (for all). Statistical comparisons for panels (**d, e**) were performed using Kruskal-Wallis test, comparing each condition to the “no cells” control. Statistics for (**d**) from left to right: p = 0.0285, 0.001, z = 2.191, 3.2386. Statistics for (**e**): p = 0.0026, z = 3.012. Significance levels: *\*p<0.05, **p<0.01, ***p<0.001, ****p<0.0001*; nonsignificant data points are not marked. Error bars in (**c**) represent mean ± s.e.m. of N = 3 biological replicates. N = *3* replicates from independent dilution series for (**d**) and (**e**). Each data point is the arithmetic mean of N = 2 technical replicates. All ultrasound image scale bars represent 1 mm.

Triply transduced and sorted cells in the MOI 60 - 4:1:1 condition showed significant BURST contrast at concentrations as low as 1e6 cells per milliliter (**Fig. 2d**). xAM was able to detect as few as 1 × 10^7^ cells per milliliter (**Fig. 2e**).

We measured the viability of Jurkat cell lines transduced with the 3-vector system, with and without inducing GV expression. Using 7-AAD (7-aminoactinomycin D) live/dead staining, we found no significant Dox-induced cell death attributed to GV expression (**Extended Data Fig. 4d-e**). However, we saw overall more cell death in cells transduced with the three vectors at MOI 60 compared to MOI 30. We attribute this to the high viral dose, which likely included a large number of empty capsids from packaging larger payloads^19^.

These experiments together demonstrated the adaptability of our lentiviral architecture for expressing GVs in immune cells, and more generally, in cells considered to be more difficult to transduce.

### Acoustic reporter genes enable monitoring of T cell activation

Having engineered immune cells to express GVs, we hypothesized that this expression could be connected to an activity-dependent promoter as a read-out of cellular state. In particular, we sought to trigger ultrasound contrast in T cells when they become activated by antigen engagement. To achieve this goal, we connected GV expression to the NFAT promoter, which activates upon T cell receptor engagement and accompanying calcium influx^20,21^. We constructed a gene circuit in which the NFAT promoter is placed upstream of the rtTA transactivator, such that NFAT activation initiates transcription of the rtTA, which in the presence of doxycycline drives GV gene transcription (**Fig. 3a**). This architecture is beneficial because it allows potentially weak NFAT activity to drive a stronger rtTA-TRE transcription complex and temporally confines GV expression to periods when it is needed for imaging, thereby minimizing cellular burden. This architecture is also modular in the sense that only a simple swap of the promoter upstream of rtTA is needed to change the conditionality of expression for all eight GV genes and accompanying fluorophores.

**Figure 3:**
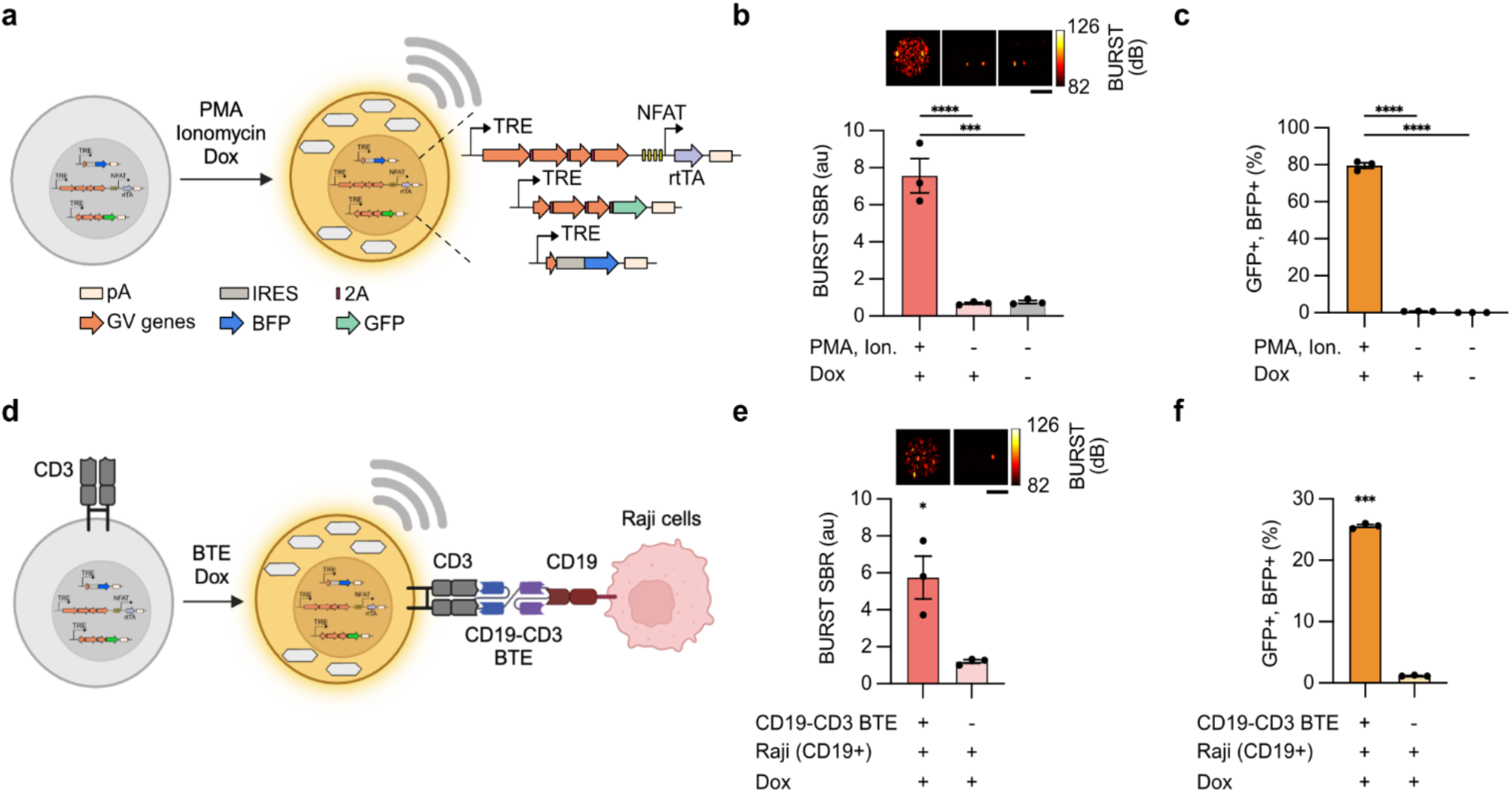
Activation-dependent GV expression enables ultrasound imaging of T cell activity. **a**, Schematic of lentiviral vectors designed to report T cell activity state by expressing GVs downstream of the NFAT promoter. T cell activation is chemically induced by incubating cells with PMA and ionomycin, which trigger the NFAT promoter to express rtTA, and in the presence of doxycycline, activate transcription of GV genes. **b**, Representative BURST images (top) and signal quantification (bottom) of Jurkat cells transduced with the vectors in panel (**a**) with or without chemical activation, normalized to wild-type (WT) cells (not shown). **c**, Percentage of Jurkat cells in panel (**b**) expressing both fluorescent reporters (GFP and BFP), with and without activation. **d**, Imaging T cell activation in response to CD19-CD3 BTE-mediated engagement with CD19+ Raji cells, using cells engineered with the vectors shown in (**a**). **e**, Representative BURST images (top) and signal quantification (bottom) of Jurkat cells engineered to express GVs upon receptor-mediated activation in the presence of Dox. **f**, Percentage of cells from panel (**e**) expressing both fluorescent reporters (GFP and BFP), with and without receptor engagement. Statistical comparisons for (**b, c**) were performed using Fisher’s LSD test, comparing the PMA, Ion+ and Dox+ condition against the other two conditions. (**b**) from left to right: p = <0.0001, 0.0001; t = 9.116, 9.028. df = 6, 6. (**c**) from left to right: p = <0.0001, <0.0001; t = 65.96, 66.60. df = 6, 6. Statistical comparisons for (**e, f**) were performed using ratio paired two-tailed t tests. (**e**): p = 0.0322, t = 5.434, df =2. (**f**): p = 0.0002, t = 67.34, df =2. Significance levels: *\*p<0.05, **p<0.01, ***p<0.001, ****p<0.0001*. Error bars represent mean ± s.e.m. of N = 3 biological replicates. Each data point represents the arithmetic mean of N = 2 technical replicates. Scale bars for all ultrasound images represent 1 mm. Ion.: ionomycin.

We created a Jurkat T cell line transduced with this engineered three-virus combination and tested whether we can turn on GV expression by chemically inducing the NFAT promoter. When we stimulated these cells with phorbol 12-myristate 13-acetate (PMA) and ionomycin^22^ in the presence of doxycycline, they produced strong ultrasound contrast (**Fig. 3b**). The signal was activity-specific, with a fold-change of 10.9 in BURST SBR when compared with doxycycline alone. Flow cytometry confirmed specific circuit activation, with an activation-dependent fold change of 99.0 in the fraction of cells expressing GFP and BFP (**Fig. 3c**).

Having confirmed the circuit design, we tested if receptor-induced activity could be detected with ultrasound when T cells engage with an on-target bait cell. We used a CD19-CD3 bispecific T-cell engager (BTE), which causes T cell activation when a BTE molecule binds CD3 on the T cell surface and CD19 on the target cell surface at the same time^23^. We co-cultured our NFAT-inducible Jurkat cell line with CD19+ Raji cells at a 1:1 ratio (**Fig. 3d**). Within 24 hours, in the presence of the BTE, we observed a large increase of ultrasound contrast (4.8-fold change in BURST SBR) relative to the same co-culture without the molecular engager (**Fig. 3e**). Flow cytometry corroborated this activation by showing a 22.1-fold change in the GFP and BFP positive population fraction (**Fig. 3f**). These results highlight the potential of ultrasound to detect cellular activation states by leveraging activity-dependent transcription to drive GV expression.

### Ultrasound monitors in vivo homing of GV-expressing therapeutic T cells into tumors

A major challenge in cell-based therapies is to monitor the trafficking of engineered cells to their intended target tissues after *in vivo* administration. To address this challenge, we endeavored to show that we could use our viral GV expression platform to label a primary T cell therapy and image its homing to tumors following systemic infusion.

To achieve this goal, we isolated primary T cells from human peripheral blood mononuclear cells (PBMCs) and transduced them with our Dox-dependent 3-vector system. Since at a total viral titer of MOI 120 (based on functional tittering in HEK cells), less than 20% of these hard-to-transduce cells^24,25^ were triple-positive for all three viruses, we used a 1:1:1 ratio of the *gvpA* to AF1 and AF2 vectors to maximize the number of cells with at least one functional unit of each construct (**Fig. 4a**, top row). After sorting the cells based on GFP expression to ensure that they received all assembly factor genes, and inducing them with doxycycline for 72 hours, we found a consistent BURST signal in Dox-induced T cells of five different human donors that was 3.8 ± 0.2 times (geometric mean of ratios ± s.e.m. of log(ratios)) higher than the signal from wild-type non-transduced cells (**Fig. 4b**), marking the first demonstration of primary mammalian cells expressing acoustic reporter genes. Of the Dox-induced sorted cells, 55.8 ± 7.36% (N = 5, mean ± s.e.m) were both GFP and BFP positive, indicating the presence of both AFs and *gvpA* in most of this population (**Extended Data Fig. 5**).

**Figure 4:**
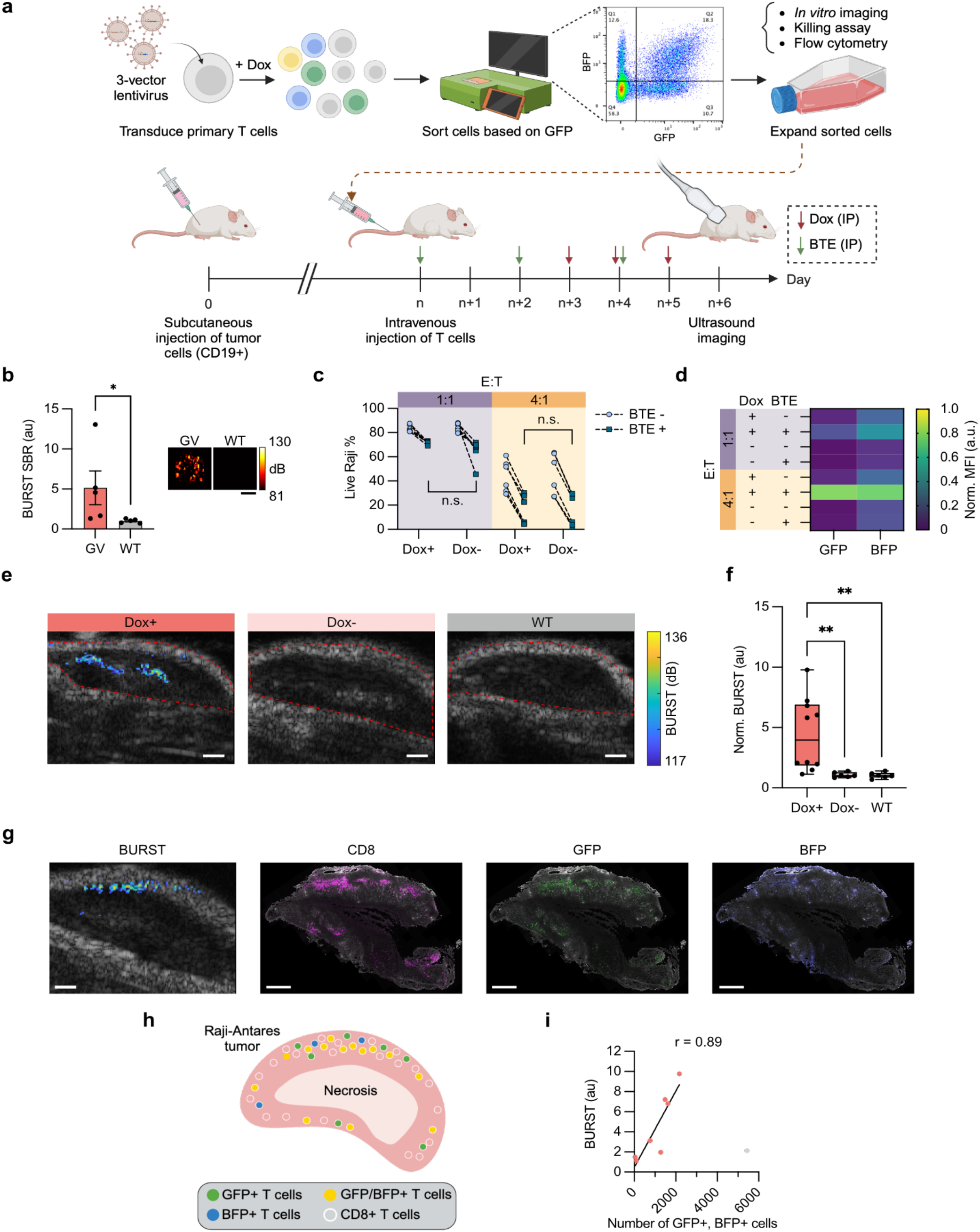
In vivo tracking and functional assessment of primary T cells engineered to express GVs. **a**, Workflow for engineering T cells isolated from human PBMCs to express Dox-inducible GVs and imaging BTE-mediated homing of cytotoxic T cells into target tumors. Top: T cells are transduced with the 3-vector lentiviral system, and GFP expressing cells are sorted to select for transduction of the assembly factor vectors. The cells are then expanded in vitro for downstream characterization and in vivo injections. Bottom: T cells are systemically administered to immunocompromised mice bearing subcutaneous CD19+ Raji cell tumors, accompanied by intraperitoneal (IP) injections of BTE every 48 hours and IP injections of doxycycline for 72 hours prior to ultrasound imaging. **b**, BURST SBR (left) and representative images (right) of GV-expressing and WT T cells. Paired one-tailed t test, p = 0.0115, t = 3.592, df = 4. Error bars represent mean ± s.e.m. of N = 5 PBMC donors; each data point is the arithmetic mean of N = 3 technical replicates. **c**, Percentage of live Raji cells in a cytotoxicity assay, where T cells are co-cultured with CD19+ Raji cells at effector-to-target (E:T) ratios of 1:1 and 4:1. Welch’s two-tailed t-test, p (from left to right) = 0.1535, 0.9973, t (from left to right) = 1.668, 0.0034, df (from left to right) = 5.244, 9.935, N = 6 biological replicates (2 PBMC donors and 3 replicates per donor). **d**, GFP and BFP fluorescence measurements from (**c**), normalized to the maximum mean fluorescence intensity (MFI) of each reporter. **e**, BURST images overlaid on anatomical B-mode grayscale images of representative Raji tumors in mice injected with GV-expressing T cells, either Dox-induced or uninduced, and mice injected with WT T cells. Red lines indicate tumor boundary used for signal quantification. **f**, Quantification of BURST signal inside tumors. Center line, median; box limits, quartiles; whiskers, min to max. Unpaired Welch’s two-tailed t test, Dox+ vs. Dox-, p = 0.0067, t = 3.483, df = 9.149; Dox+ vs. WT, p = 0.0063, t = 3.525, df = 9.194. N = 10 mice for Dox+, N = 6 for Dox-, N = 6 for WT. **g**, Histology of an example tumor infiltrated by GV-expressing T cells and its corresponding BURST image. White: Raji-Antares cells; Magenta: CD8+ cytotoxic T cells; Green: GFP+ T cells; Blue: BFP+ T cells. **h**, Schematic of T cell distribution in the Dox-induced tumor from (**g**). **i**, Correlation of the BURST signal within tumors with the absolute numbers of GFP+ and BFP+ T cells in corresponding histological sections. Pearson correlation: r = 0.89, p = 0.0072 (two-tailed), N = 7 mice. One mouse (gray data point, mouse 3 in **Extended Data** Fig. 7a) was excluded due to a significant number of T cells located outside the imaging plane whose GV expression was not captured by BURST (see **Supplementary Fig. 1**). All scale bars represent 1 mm.

After demonstrating ultrasound contrast, we asked if GV expression affects T cell cytotoxicity, a critical function for administered therapies. To investigate this, we co-cultured the T cells with CD19+ Raji cells at 1:1 and 4:1 effector-to-target (E:T) ratios in the presence of the CD19-CD3 BTE, and measured the percentage of live target cells with 7-AAD staining 24 hours after co-culture. We found that GV expression (controlled with doxycycline induction) did not affect T cell cytotoxicity at either ratio, and as expected the cells were more effective at killing at the higher E:T ratio and in the presence of the targeting BTE (**Fig. 4c**). Interestingly, flow cytometry showed increased GFP and BFP expression in co-cultured T cells with BTE engagement, especially at the higher E:T ratio (**Fig. 4d**). This is because increasing the E:T ratio leads to more efficient target cell lysis, and hence increased T cell activity and protein expression^26–29^.

We then examined whether GV expression in primary T cells affects their activation state. To assess this, we used well-established phenotypic and functional readouts that capture key aspects of T cell activation and effector differentiation. Specifically, we performed surface staining for three activation markers: CD69, an early activation marker rapidly upregulated following TCR stimulation^30^, and CD38 and HLA-DR which are associated with sustained activation and effector differentiation^31^. After 3 days of Dox induction, we found no difference in CD69 and HLA-DR expression levels in Dox-induced and uninduced cells, and only a small difference in CD38 expression levels of induced GV expressing cells (**Extended Data Fig. 6a-c**). We also measured secretion of IFN-γ, a canonical effector cytokine produced by activated T cells^32^, in the culture supernatant and observed no significant difference between Dox-induced and uninduced GV expressing T cells, and a small difference in Dox-induced wild-type T cells, likely due to effects of co-culturing with doxycycline (**Extended Data Fig. 6d**). Lenti-transduced cells showed overall reduced HLA-DR expression and IFN-γ release compared to wild-type untransduced T cells, likely reflecting effects of viral transduction.

To determine whether BURST imaging impacts T cell function or viability, we Dox-induced T cells for 3 days and exposed them to the BURST pulse sequence while in suspension. We then cultured them for an additional 36 hours and assessed activation marker expression and viability by flow cytometry (**Extended Data Fig. 7a**). We found no significant differences in activation state between BURST-exposed and unexposed cells (**Extended Data Fig. 7b-d**), and cell viability was not affected (**Extended Data Fig. 7e**).

Having demonstrated successful GV expression and ultrasound imaging *in vitro*, we designed an experiment to show that primary T cells could be imaged *in vivo* after systemic administration, allowing ultrasound to visualize their trafficking to tumors. We intravenously infused the engineered T cells into immunocompromised mice bearing subcutaneous CD19+ Raji cell flank tumors, followed by intraperitoneal (IP) injections of the CD19-CD3 BTE every 2 days to stimulate activation and local expansion inside the tumor microenvironment^23,33^. On day 3 after T cell infusion, we induced GV expression with intraperitoneal doxycycline and imaged the tumors with ultrasound (protocol depicted in **Fig. 4a**, bottom row).

Ultrasound imaging revealed clear BURST ultrasound contrast in the tumors of mice infused with GV-expressing T cells and induced with doxycycline (**Fig. 4, e-f**). This signal was absent in mice that were not Dox-induced or received wild-type T cells. This response was consistent across our total cohort of 22 mice, with 4.1 fold-change in BURST SBR for the Dox+ vs. Dox-groups, and 4.3 for the Dox+ vs. WT groups (**Fig. 4f** and **Extended Data Fig 8**). In a subset of animals, we collected tumors and performed immunofluorescence imaging to confirm that the localization of ultrasound signal observed in tumors matches the spatial distribution of GV-expressing T cells observed by histology, finding a clear correspondence (**Fig. 4g** and **Extended Data Fig. 9a**). For example, in a representative tumor, the core was mostly necrotic with few live cells, while both ultrasound and histology showed cytotoxic T cells expressing our reporter construct concentrated at the viable tumor periphery near the top of the tumor (**Fig. 4, g-h**). Additionally, flow cytometry analysis of digested tumors showed that 14.9 ± 7.8% of cytotoxic CD8+ T cells (N = 5 mice) were GFP+/BFP+, indicating successful co-expression of all transgenes (**Extended Data Fig. 9b**). While this fraction could be further increased through stricter sorting or additional enrichment strategies, it was sufficient to generate detectable ultrasound signal *in vivo*.

Consistent with the pressure-dependent nature of BURST imaging, signal was strongest near the primary acoustic focus, where pressure is maximal. At depths above and below this focal region, pressure may still be sufficient to collapse some GVs but insufficient to generate strong detectable contrast, which can reduce signal from those regions when subsequently imaged (**Supplementary Fig. 1**).

The ability of ultrasound to visualize this sub-tumor localization *in vivo* is in stark contrast to the diffuse signal typically obtained with bioluminescence, and complements nuclear imaging approaches that can provide whole-body sensitivity but often with lower spatial resolution in comparable settings. Notably, tumors with low ultrasound contrast also had few T cells by histology (**Extended Data Fig. 9a**). Indeed, we found a strong positive correlation between the absolute number of GFP+ and BFP+ T cells counted in a given histology slice *postmortem* and the BURST signal measured in the corresponding ultrasound image plane *in vivo* (r = 0.89, p = 0.0072, Pearson correlation, **Fig 4i**), suggesting that ultrasound can provide a quantitative noninvasive readout of cell therapy homing and/or expansion. We note that variability in BURST signal across mice likely reflects biological heterogeneity, potentially driven by differences in T cell infiltration and expansion within tumors, as corroborated by histology.

Overall, these experiments demonstrated, for the first time, an ultrasound method to noninvasively visualize the homing of a mammalian cell therapy into its target tissue.

### Intratumoral ultrasound signal from GV-expressing CAR-T cells correlates with therapeutic efficacy

We next asked whether our ultrasound imaging approach could predict the therapeutic efficacy of chimeric antigen receptor (CAR) T cell therapies in solid tumors. We engineered primary T cells to co-express the GV gene cluster together with an anti-CD19 CAR. We incorporated the gene encoding CAR into the *gvpA* vector of our three-lentivirus system (**Fig. 5a**), ensuring that cells with the therapeutic gene, would also retain the main structural gene *gvpA* required for forming complete GVs.

**Figure 5:**
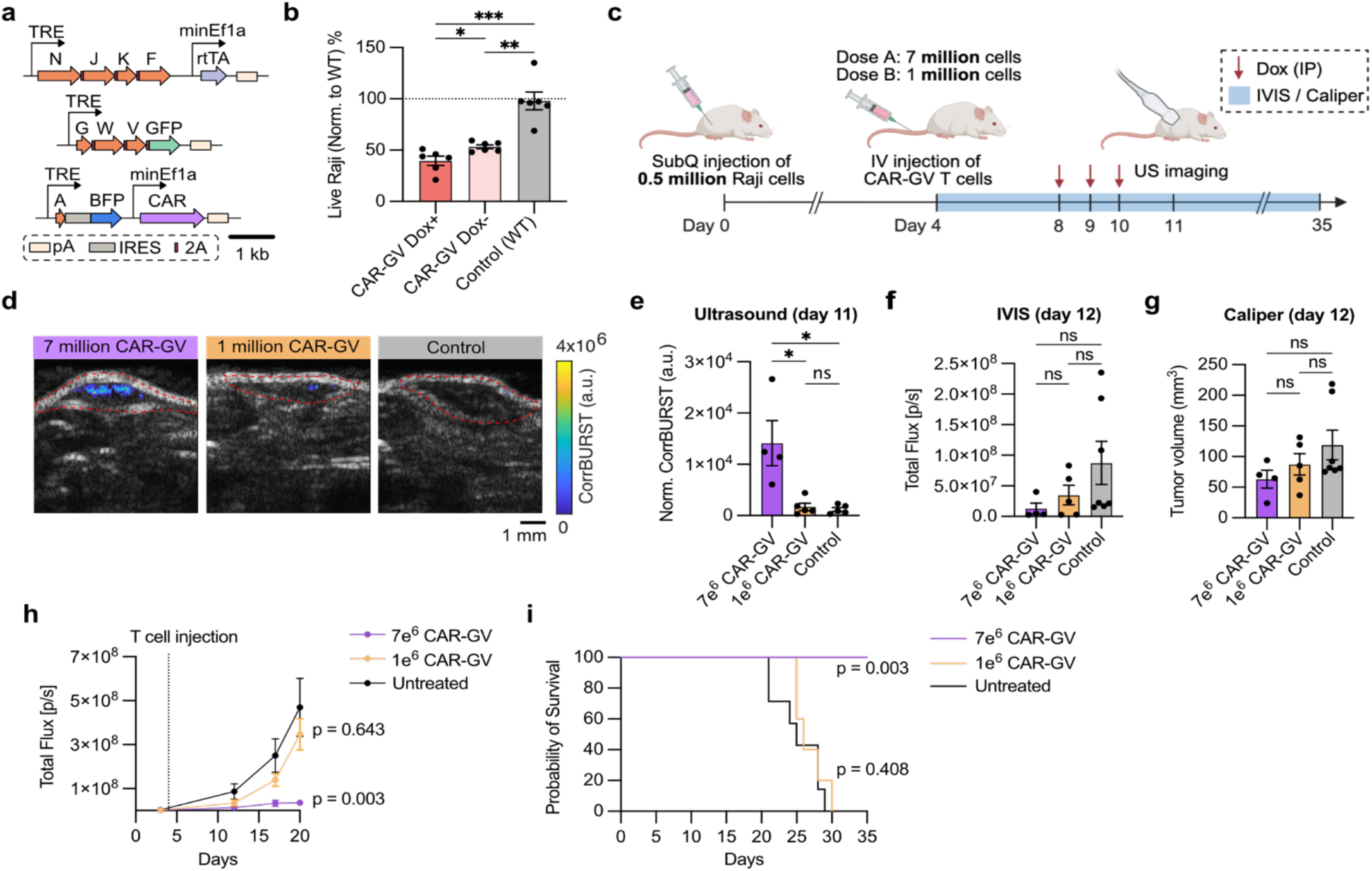
Ultrasound imaging of CAR-GV T cells quantitatively measures intratumoral homing. **a**, Genetic design of a 3-vector lentiviral system encoding a CD19 CAR and GV genes (CAR-GV). **b**, In vitro killing efficiency of CAR-GV T cells with and without Dox induction, compared to untransduced (WT) T cells at E:T ratio of 5:1; n = 2 biological replicates from different donors with n = 3 technical replicates per donor. One-tailed Welch’s t test. p (Dox+ vs. Dox-) = 0.0134, p (Dox+ vs. WT) = 0.002, p (Dox- vs. WT) = 0.0015. **c,** Experimental timeline for in vivo assessment of anti-tumoral therapeutic efficacy and ultrasound imaging of CAR-GV T cells in NSG mice bearing subcutaneous Raji-Antares tumors. **d,** Representative day 11 CorrBURST image (colorbar) overlaid on anatomical B-mode image (grayscale) of Raji tumors in mice injected with Dose A (7e^6^) or Dose B (1e^6^) CAR-GV T cells, or non-GV–expressing T cells (Control); red lines indicate tumor boundaries used for signal quantification. **e,** Sum of CorrBURST signal intensities within the entire tumor volume, normalized to tumor size (Norm. CorrBURST). 7e^6^ CAR-GV, N = 4; 1e^6^ CAR-GV, N = 5; Control, N = 5 mice. Two-tailed Mann-Whitney test. p (7e^6^ vs. 1e^6^) = 0.016, p (7e^6^ vs. Control) = 0.016, p (1e^6^ vs. Control) = 0.841. **f,** Total flux from tumors measured by IVIS on day 12. Two-tailed Mann-Whitney test. p (7e^6^ vs. 1e^6^) = 0.730, p (7e^6^ vs. Control) = 0.073, p (1e^6^ vs. Control) = 0.432. **g,** Tumor size measurements on day 12. Two-tailed Mann-Whitney test. p (7e^6^ vs. 1e^6^) = 0.556, p (7e^6^ vs. Control) = 0.109, p (1e^6^ vs. Control) = 0.639. **h,** Total flux of tumors measured by IVIS over 35 days following tumor engraftment. 2-way ANOVA test comparing treatment groups to Control group. **i,** Kaplan–Meier survival probability of the three experimental groups. Long-rank (Mantel-Cox) test comparing treatment groups to Control. **(e, f, g, h, i)** 7e^6^ CAR-GV, N = 4; 1e^6^ CAR-GV, N = 5. Control, N=7. Significance levels: ns p≥0.05, *p<0.05, **p<0.01, ***p<0.001, ****p<0.0001. Error bars represent mean ± s.e.m.

We first tested whether GV expression affects cytotoxic function of CAR-T cells. Primary T cells were transduced with all three GV lentiviral vectors and sorted based on fluorescence expression to obtain CAR-GV T cells. These cells were co-cultured with CD19+ Raji-Antares target cells at an effector-to-target ratio of 5:1 for 24 hours, with or without Dox induction of GV expression. Flow cytometry showed that only 39.64 ± 4.46% (mean ± s.e.m) of Raji cells remained viable after co-culture with Dox-induced CAR-GV T cells, compared with 53.30 ± 1.92% (mean ± s.e.m) viability in the uninduced CAR-GV co-culture (**Fig. 5b**). These results indicate that GV expression does not impair CAR-T cytotoxic activity.

We next confirmed that CAR-GV T cells could be detected *in vivo* using ultrasound. CAR-GV T cells were intravenously injected into mice bearing subcutaneous Raji tumors, and GV expression was induced with Dox for 3 days prior to imaging on day 7. Dox-induced mice displayed strong BURST contrast within their tumors, whereas uninduced mice or mice receiving wild-type T cells showed minimal signal (**Extended Data Fig. 10a, b**). These results demonstrate that ultrasound can detect CAR-T cell trafficking and engraftment in solid tumors following systemic administration.

To assess whether we could track CAR-T cells over time, we performed longitudinal imaging of mice treated with CAR-GV T cells under subtherapeutic conditions, ensuring that tumors persisted during this extended imaging timeframe. BURST signal within tumors was comparable between days 7 and 21 after T cell infusion in Dox-induced mice (**Extended Data Fig. 10c-e**), indicating sustained presence of CAR-GV T cells within tumors. In contrast, mice receiving wild-type T cells exhibited low signal on day 7 and were euthanized prior to later imaging time points due to tumor burden, precluding longitudinal comparison. Together, these results show that GV-expressing CAR-T cells can be re-imaged at least two weeks after an initial BURST acquisition, supporting the feasibility of longitudinal monitoring on a therapeutically relevant timescale^34,35^, although the minimum interval required for signal recovery remains to be determined.

Having established *in vivo* detectability, we asked whether early ultrasound measurements could predict therapeutic response. Conventional assessments of CAR-T efficacy in animal models rely on changes in tumor size measured by calipers or optical imaging, which often lag behind cellular events occurring within the tumor. To investigate whether ultrasound provides an earlier indicator of response, we injected mice bearing subcutaneous Raji-Antares tumors with either a high dose (7 × 10^6^) or sub-therapeutic low dose (1 × 10^6^) of CAR-GV T cells 4 days after tumor implantation. GV expression was induced with Dox for three days prior to ultrasound imaging (day 11). Tumor growth was then monitored longitudinally using caliper measurements and luminescence imaging until day 35 or humane endpoint (**Fig. 5c**).

BURST imaging on day 11 revealed a clear difference in intratumoral signal between mice treated with the high and low CAR-T doses (**Fig. 5d** and **Extended Data Fig. 11**). Notably, ultrasound detected significantly higher GV-specific signal in mice receiving 7 × 10^6^ CAR-GV T cells, at a timepoint when neither caliper nor luminescence measurements via IVIS imaging showed significant differences in tumor sizes between the three groups (**Fig. 5e-g**).

Long-term monitoring revealed that tumors treated with the high CAR-T dose underwent substantial regression over the following weeks, with 1 of 4 tumors completely resolving (**Fig. 5h** and **Extended Data Fig. 12**). In contrast, tumors treated with the low CAR-T dose or left untreated continued to grow. The early difference in ultrasound signal therefore preceded and predicted the subsequent therapeutic response. Together, these results suggest that ultrasound imaging of GV-expressing CAR-T cells provides an early, non-invasive indicator of treatment efficacy.

### Immunogenicity prediction for acoustic reporter genes

The potential immunogenicity of reporter genes is an important consideration for future clinical translation. To obtain an initial assessment of GV protein immunogenicity, we performed in silico HLA class I antigen-presentation analyses using two widely used prediction tools, MUNIS^36^ and NetMHC4.1^37^. As a benchmark, we compared GV proteins with HSV-TK, a reporter gene commonly used in preclinical studies^38^, GFP and mScarlet-I, and the bacterial nucleases SaCas9 and AsCas12a, both of which are being translated into clinical use. Both prediction tools produced similar rankings of candidate epitopes across the GV gene cluster (**Extended Data Fig. 13**), sharing multiple predicted epitopes (**Supplementary Table 2**). The largest GV protein, *gvpN*, contained the highest number of predicted presented peptides, whereas most other GV proteins exhibited lower predicted epitope numbers. The predicted number of epitopes for individual GV proteins was generally comparable to or lower than those of HSV-TK. Moreover, the cumulative predicted burden of the full GV gene cluster was comparable to or lower than that of SaCas9 and AsCas12a, for which immunogenic epitopes have been experimentally verified and rationally redesigned to reduce MHC binding and CD8+ T cell recognition while preserving protein function^39^. This analysis suggests that GV expression may elicit immune recognition in humans, consistent with the non-human origin of GV proteins, similar to other reporter genes that are widely used in preclinical research and genome-engineering proteins shown amenable to deimmunization and clinical development.

## DISCUSSION

This work introduces a genetic technology platform enabling the use of acoustic reporter genes to image a broader variety of cell types, including primary cells deployed in an *in vivo* cell therapy context. By devising a set of suitable viral constructs, we make it possible to introduce GV genes into transfection-resistant cells such as T cells, with our most effective system comprising a combination of three lentiviral vectors encoding the 8 essential GV genes, along with an rtTA transactivator and fluorescent proteins for correlative histology. In addition, we showed how this expression system can be easily modified to monitor cellular states using activity-dependent promoters. Furthermore, we demonstrated that this system enables the noninvasive *in vivo* monitoring of therapeutic T cell homing and proliferation in tumors with a combination of penetration depth and resolution not available with other modalities. Importantly, we extend this capability beyond tracking by showing that intratumoral ultrasound signal from GV-expressing CAR-T cells correlates with and predicts therapeutic efficacy at early time points, before conventional measurements detect treatment response.

This acoustic imaging toolkit will significantly advance preclinical studies by allowing the study of the dynamics of cell-based therapies *in vivo*, such as how adoptively transferred T cells infiltrate solid tumors and where they expand within the tumor microenvironment. Additionally, it could facilitate the monitoring of off-tumor on-target cytotoxicity, a common side effect of CAR-T cells by imaging bystander organs. More broadly, our approach opens new possibilities to study the *in vivo* dynamics of other cell-based therapies, including those based on natural killer (NK) cells, macrophages, stem cells, and any other cell types that can be engineered with viruses, and to track cellular states beyond T cell activation, such as during different stages of development.

As expected for the first demonstration of a new imaging approach, our toolkit can benefit from further improvements. The high MOI required for efficient co-transduction of multiple vectors in primary T cells^40^ could be burdensome for certain cell types and expensive to manufacture at scale. We anticipate that this limitation will be overcome with future improvements in GV expression efficiency through modified assembly factor stoichiometry^10^, mutagenesis of the GV genes^41^, or alternative delivery methods strategies such as mRNA or lipid nanoparticle based approaches^42^. Further gains could be made by optimizing the expression cassette with improved cell type-specific promoters, targeted genomic integration and addition of elements to reduce epigenetic silencing^43^.

Another limitation is that our primary T cells were detectable only with collapse-based BURST imaging, which, while highly sensitive, requires re-expression of GVs for repeated imaging. This is likely because the GVs formed in primary T cells are too small to undergo the reversible mechanical buckling needed for non-destructive xAM imaging^44,45^. The high sensitivity of BURST can also lead to low-level background signal from small gas inclusions or other nonlinear scatterers in the imaging volume, analogous to background fluorescence in optical imaging. Future improvements in GV engineering that enhance non-destructive contrast (e.g., via xAM) would permit repeated imaging without collapse and may reduce susceptibility to such background signal. Additionally, using more advanced ultrasound hardware, such as matrix or row-column arrays, could improve the sensitivity and fidelity of xAM^46^ and BURST imaging by generating a more homogeneous pressure field within the target tissue and minimizing motion artifacts.

Beyond preclinical research, we envision the use of acoustic reporter genes in future clinical applications, allowing physicians to check that the therapy they administer reaches the right tissues and performs their intended activities. For example, after being administered a CAR-T cell therapy targeting a solid tumor, a patient could return to the clinic for imaging to ensure that the CAR-T cells have populated one or more tumor masses. In this setting, the reporter genes could be expressed transiently or induced with small molecule drugs just before imaging, thereby minimizing expression burden. Such monitoring would benefit from the ubiquitous accessibility of ultrasound imaging devices, and in the future could even be continuous using wearable bio-adhesive ultrasound patches^47–50^.

To achieve this milestone, the clinical translation of acoustic reporter genes will require a deeper understanding of their immunogenicity in immunocompetent hosts. Although our computational analyses suggest a predicted epitope burden comparable to several established reporter genes commonly used in preclinical research, future studies are needed to define the host response to GV expression *in vivo*. In CAR-T cell monitoring applications, several factors may mitigate this concern: First, GV expression could be restricted to short imaging windows after lymphodepletion, when host responses to foreign antigens may be attenuated^51^. Second, genetically labeling a relatively small subset of administered cells is sufficient to visualize population behavior (14.9 ± 7.8%, **Extended Data Fig. 9b**). Third, GV proteins, which derive from environmental bacteria or archaea, may be less likely than HSV-TK or commonly used Cas nucleases to encounter pre-existing adaptive immunity^38,52^. Finally, emerging rational and machine learning-guided deimmunization approaches ^39,53,54^, together with further minimization of the GV gene cluster^55^, may provide a route to reducing immunogenic epitopes while preserving GV function.

Using the technology developed in this study, ultrasound can now be utilized to image primary mammalian cells, including therapeutic cells, *in vivo*. With further improvements, getting a quick ultrasound readout may someday become as common for assessing the status of cell therapies as it is for routine anatomical examination.

## METHODS

### Plasmid construction and molecular biology

All constructs were made via KLD mutagenesis or Gibson Assembly using enzymes from New England BioLabs. All plasmids and their sources of genetic material are described in **Supplementary Table 1**. Constructs were cloned in NEB Turbo Competent *E. coli* (High Efficiency, Catalog #C2984). Fluorescent reporters referred to in the text as GFP and BFP are mEGFP and mTagBFP2, respectively.

### Lentivirus packaging, titrating, and transduction

#### Packaging

DMEM+ media for packaging was prepared by supplementing high-glucose DMEM + GlutaMAX containing 1 mM sodium pyruvate (Gibco, Catalog #10569), with 20 mM HEPES (Cytvia, Catalog #SH30237), 1X MEM-Non-Essential Amino Acids (0.1 mM, Gibco #11140), and 10% fetal bovine serum (bio-techne, Catalog # S10350). HEK293T cells (ATCC, CRL-3216), passaged no more than 20 times, were seeded in 10 cm plates and grown in DMEM+ until they reached 80 - 90% confluence. A mixture of 22 µg pLV (plasmid with the gene of interest), 22 µg pCMVR8.74, and 4.5 µg pMD2.G was prepared. These plasmids were combined with polyethylenimine (PEI Max, Polysciences) at a PEI to DNA mass ratio of 3:1, incubated at room temperature for 12 minutes, and then added to the cells. Cells were incubated at 37°C and 5% CO2 for 12 hours. After this period, the transfection media was replaced with 10 mL of fresh DMEM+. After 48 hours, the spent media was collected and centrifuged at 500g for 10 minutes to remove debris. The viral supernatant was then concentrated 100-fold into 1X phosphate-buffered saline (PBS) using LentiX Concentrator (Takara Bio, Catalog #631232), following the manufacturer’s protocol. The concentrated virus was aliquoted and stored at −80°C for up to 6 months, avoiding freeze-thaw cycles.

#### Measuring titer

For the NFAT-inducible virus, LentiX GoStix Plus (Takara Bio, Catalog #631281) was used to measure the titer, following manufacturer’s instructions. All other viruses were functionally titrated in various HEK-based cell lines, as described below, following the protocol outlined here. Cells were seeded in 24-well plates in DMEM and grown to 70 - 90% confluence. Three wells were trypsinized, and the average of their cell count was used in calculating the titer, described below. A serial dilution of the virus was prepared in fresh DMEM and added to each well. Plates were spun for 50 minutes at 800g at 35°C and then incubated at 37°C and 5% CO2 for 16-20 hours. The media was aspirated and replaced with fresh DMEM, supplemented with doxycycline (1 µg/mL) for the HEK-TetON and HEK-TRE cell lines. Cells were incubated for an additional 24 hours before being prepared for flow cytometry. Gating strategy for flow cytometry is shown in **Supplementary Fig. 2**. The functional titer of each virus was calculated using the following formula:

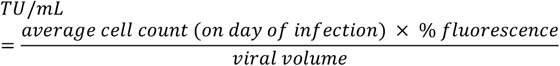

Only dilutions resulting in less than 30% fluorescent cells were used for the calculation. This threshold was chosen because it ensures that the probability of any cell receiving more than one functional viral particle is less than 26%.

The following HEK293T-based cell lines were used for functional titering: 1) wild-type HEK293T cells used to titrate any virus expressing a fluorescent protein constitutively, 2) HEK-TetON cells used to titrate any virus expressing a fluorescent protein downstream of the TRE promoter, 3) HEK-TRE cells engineered to express mScarlet-I downstream of TRE, and used to titrate any virus expressing rtTA.

#### Transduction

To transduce HEK293T cells, the cells were seeded in 12-well plates 24 hours before transduction and grown to 70 - 90% confluence. On the day of transduction, cells from three wells were counted and averaged to determine the viral volume needed for each desired MOI. Lentivirus was mixed with fresh media and added to each well with 10 µg/mL of polybrene (Millipore Sigma, Catalog #TR1003). Cell culture plates were spun at 800g for 50 minutes at 35°C, and incubated at 37°C for 16-20 hours. The viral media was then removed, and the cells were passaged for downstream applications.

To transduce suspension cells (Jurkats and primary T cells), the cells were counted on the day of transduction and one million cells were mixed with fresh media and virus at the desired MOI. The cells were spun at 1000g for 50 minutes at 35°C, then incubated at 37°C for 16-20 hours. Viral media was then removed, and the cells were passaged for further use.

### T cell viability and surface staining

Cell viability was assessed using 7-AAD (BioLegend, Cat. #420403) according to the manufacturer’s protocol. Briefly, cell pellets were resuspended in 1000200 µL of 1X PBS, and 7-AAD was added at a 1:100 dilution. Cells were incubated for 5-10 min in the dark and analyzed immediately by flow cytometry using a MacsQuant Analyzer 10.

For cell surface staining, cells were pelleted and resuspended in antibody dilution buffer (0.5% BSA in 1X PBS). Antibodies were added at manufacturer-recommended dilutions, and cells were incubated for 30 min at 4 °C in the dark. Cells were then washed twice and resuspended in 1X PBS prior to flow cytometry analysis.

Surface staining of activation markers was performed using the following antibodies: CD69 (BioLegend, 310910), CD38 (Miltenyi, 130-113-429), and HLA-DR (Miltenyi, 130-111-790).

### IFN-γ release assay

IFN-γ secretion was measured following a previously established protocol^56^ with minor modifications. Briefly, on day 0 (day 8 post T cell isolation and stimulation), T cells were seeded in 96-well plates and cultured in the presence or absence of doxycycline, with IL-2 added to all conditions. Doxycycline was maintained daily through day 3. On day 4, cells were split into triplicate wells and cultured in the presence of IL-2 until day 8 without further manipulation. Cells were then stimulated with anti-CD3 antibody (Thermo, 13-0037-82) for 4 hours, after which supernatants were collected for analysis. IFN-γ levels were quantified using a human IFN-γ ELISA kit (BioLegend, 430107) according to the manufacturer’s instructions, and absorbance was measured using a Tecan Spark plate reader.

### Cell sorting

Cells were expanded without prior doxycycline addition and induced for the first time with doxycycline (1 µg/mL) 24 hours before sorting. Cells were collected and resuspended in cell sorting buffer (Miltenyi Biotec, Catalog #130-107-207) at a concentration of 2-4 million cells/mL, and GFP- and BFP-expressing cells were sorted using the MACSQuant Tyto cell sorter. For primary T cells, doxycycline induction (1 µg/mL) was done for 48 hours, followed by resuspension in RPMI media without FBS, and sorting based on GFP expression. Sorted cells were immediately resuspended in their respective media and incubated at 37°C, with a recovery period of at least 24 hours before use in downstream experiments.

### In vitro T cell activity induction

For chemical induction, Jurkat cells (Clone E6-1, ATCC #TIB-152) were stimulated with 50 ng/mL phorbol 12-myristate 13-acetate (VWR, Catalog #16561-29-8) and 1.3 µM ionomycin (Sigma, Catalog #I9657) for 24 hours before being used for flow cytometry (MacsQuant Analyzer 10) and ultrasound imaging. For receptor-mediated induction, Jurkat cells were co-cultured with Raji cells (ATCC, CCL-86) at a 1:1 ratio in media supplemented with 1 ng/mL CD19-CD3 BTE (BPS Bioscience, Catalog #100441). Similar to chemical induction experiments, cells were used for flow cytometry and ultrasound imaging after 24 hours of co-culture.

### T cell isolation, transduction, and sorting

T cells were isolated from frozen human peripheral blood mononuclear cells (STEMCELL, Catalog #70025.1) using the EasySep Human T Cell Isolation Kit (STEMCELL, Catalog #17951), and activated with Dynabeads Human T-Activator CD3/CD28 beads (Gibco, Catalog #1131D). T cells were cultured in RPMI supplemented with GlutaMAX (Gibco, Catalog #61870036) and recombinant human IL-2 at 100 U/mL (Miltenyi Biotec, Catalog #130-097-743). 16-20 hours after isolation/activation, the cells were transduced with lentivirus as described above. Media containing virus was removed 16-20 hours after transduction. The cells were passaged every other day to maintain a density of 1e6 cells/mL, and provided with IL-2 at 100 U/mL. Activation beads were removed on day 5 post-isolation/activation. Fluorescence was induced with doxycycline for 2 days, and GFP-positive cells were sorted using the MACSQuant Tyto Cell Sorter (Miltenyi Biotec) on day 7. The sorted cells were then expanded *in vitro* for further use.

### Cell preparation and phantom loading for in vitro imaging

Adherent cells were seeded 24 hours in advance and allowed to reach 70 - 80% confluence. They were then induced with doxycycline hyclate at 1 µg/mL every 24 hours for 72 hours. On the day of imaging, cells were trypsinized, spun down at 300g for 5 minutes, and resuspended in 1X PBS. They were briefly mixed with 1% melted agarose in 1X PBS (1:1 dilution at 42°C) and loaded into agarose phantoms at a final concentration of 25 million cells/mL for all experiments, except detection limit experiments.

Suspension cells were resuspended to 0.8e6 cells/mL on the day of induction, and supplemented with doxycycline at 1 µg/mL every 24 hours for 72 hours. Meanwhile the cells were passaged to keep their density below 1e6 cells/mL. On the day of imaging, cells were spun down at 300g for 5 minutes and resuspended in 1X PBS. They were briefly mixed with 1% melted agarose in 1X PBS (1:1 dilution at 42°C) and loaded into agarose phantoms at a final concentration of 25 million cells/mL for all experiments, except detection limit experiments. Primary T cells were loaded at 50 million cells/mL.

### In vitro killing assay

Primary T cells were mixed with Raji cells to achieve a final density of 1 million cells/mL. To induce receptor-mediated T cell cytotoxicity, CD19-CD3 BTE (1 ng/mL, BPS Bioscience, #100441) and IL-2 (100 U/mL) were added. After 24 hours of co-culture, the percentage of live Raji cells was measured using 7-AAD (BioLegend, Catalog #420403) staining with flow cytometry, following the manufacturer’s protocol.

### Tumor implantation and T cell injections in mice

NSG mice (NOD-scid IL2Rgamma, JAX, Strain #005557) aged 3 to 8 months were used. For BTE experiments, 4 × 10^6^ Raji cells expressing the Antares fluorescence reporter^57^ under the Ef1a promoter were resuspended in matrigel (Corning, Catalog # 354234) on ice and injected subcutaneously near the flank. When tumors reached a measurable size between days 12 to 17 after implantation, the mice were infused with 1 × 10^7^ T cells via tail-vein injections. BTE (0.25 mg/kg in saline, BPS Bioscience, #100441) and doxycycline hyclate (4 mg/kg, in saline) were administered intraperitoneally according to the timeline in **Fig. 4a**, with the first BTE dose administered 30 min before T cell injection. Hair around the tumor was shaved immediately before imaging, and mice were anesthetized under 1-2.5% isoflurane during imaging.

For repeated imaging experiments, mice were implanted subcutaneously with 1 × 10^6^ Raji-Antares cells. On day 7 post-implantation, mice received 5 × 10^6^ CAR-GV T cells via systemic injection. Tumors were imaged by ultrasound on days 14 and 21.

For therapeutic dose experiments, mice were implanted with 5 × 10^5^ Raji-Antares cells and treated on day 4 with either 1 × 10^6^ or 7 × 10^6^ CAR-T cells. Tumors were imaged by ultrasound on day 11. Tumor progression was subsequently monitored by caliper measurements and bioluminescence imaging until day 35 or humane endpoint.

### Immunohistochemistry of tumors

Tumors were extracted from mice following ultrasound imaging and fixed overnight in 4% paraformaldehyde at 4°C with continuous rotation. They were then immersed in 30% sucrose for 48 hours before embedding in O.C.T. Compound (Fisher Scientific, Catalog #23-730-571) at −80°C. The frozen tissue was cryosectioned at a thickness of 20 µm, with 3 to 6 slices collected every 1 mm and placed directly onto microscope slides (Fisher Scientific, Catalog #12-550-19). The location of each cryosectioned slice was recorded to match it to the corresponding BURST plane with 1 mm accuracy. Slides were dried overnight at room temperature and stored at −80°C.

Before staining, slides were brought to room temperature and washed in 1X PBS for 5 minutes, repeated three times. They were then incubated in freshly prepared blocking buffer (5% bovine serum albumin in 1X PBS) for 60 minutes at room temperature. After blocking, slides were washed in 1X PBS for 5 minutes, repeated twice. Most of the PBS was then removed, and a liquid blocker pen (Ted Pella, Catalog #22311) was used to draw boundaries around tumor slices on each slide. The antibody dilution was prepared in freshly made staining buffer (0.5% bovine serum albumin in 1X PBS). To each slide, 200 µL of the antibody solution was added, ensuring it covered all tumor slices, and slides were incubated overnight at 4°C in a humid chamber. The slides were washed in 1X PBS for 5 minutes, repeated three times, and excess PBS was removed.

Next, 200 µL of mounting media (Electron Microscopy Sciences, Catalog #17985-11) was added to each slide, which was then sealed with a coverslip. Slides were allowed to dry before imaging and protected from light throughout all staining steps. The following antibody was used for staining CD8+ T cells: Alexa Fluor 647-conjugated anti-human CD8 antibody (BioLegend, Catalog #344726) at a 1:50 dilution.

Confocal imaging was performed using an inverted laser scanning confocal microscope with a 20x objective lens. Cell counting was conducted using the Imaris software’s Spots Detection function based on GFP and BFP expression. Briefly, images were initially processed with a Gaussian blurring filter. The Spots Detection function was applied separately to the GFP and BFP channels to mark fluorescent cells. The number of overlapping GFP+ and BFP+ spots inside the tumor were then counted.

### Ultrasound data acquisition and image analysis

#### In vitro imaging

Ultrasound imaging for *in vitro* experiments was performed using a Verasonics Vantage programmable ultrasound scanning system with an L22-14vX 128-element linear array transducer (Verasonics). Image acquisition was conducted using previously published BURST and xAM scripts. For BURST imaging, the transmit waveform was set to a frequency of 15.6 MHz, with a 67% intra-pulse duty cycle, and 7 half-cycle transmits. The BURST pulse sequence consisted of a single low-pressure frame (transducer voltage: 1.6 V; peak positive pressure: 0.4 MPa) followed by forty-five high-pressure frames (transducer voltage: 15 V; peak positive pressure: 3.6 MPa). The programmable transmit focus was set to 8 mm to align with the fixed elevation focus of the transducer.

All *in vitro* BURST images were reconstructed using a temporal-template unmixing algorithm across individual pixel locations in the frame stack^16^. Signal-to-background ratio (SBR) was defined as the mean BURST signal within an ROI drawn inside the well, divided by the mean BURST signal in WT/non-transduced wells for each experiment.

For xAM imaging, cells were imaged over a range of pressures (peak positive pressure 0.2 MPa to 1.1 MPa; transducer voltage 3.5 V to 9.5 V). The pressure yielding the highest signal-to-noise ratio before GV collapse was selected for quantification. xAM SBR was defined as the mean xAM signal within an ROI drawn inside the well, divided by the mean signal in a noise ROI drawn in the phantom adjacent to the well.

#### In vivo imaging

The same Verasonics Vantage system and transducer were used in both *in vitro* and *in vivo* experiments. The BURST pulse sequence began with a single low-pressure frame (transducer voltage: 1.6 V), followed by five high-pressure frames (transducer voltage: 25 V). Multiple transmit foci were employed to capture the entire tumor, with a BURST image acquired at each focus. These images were summed to create a composite BURST image. Each cross section of tumor was imaged twice, with the second acquisition used for background normalization (details below).

BURST images were reconstructed by subtracting the 5th post-collapse frame from the collapse frame to isolate collapse-based nonlinear scattering from linear scatterers. Images with different transmit foci were then summed to create a composite image, which was subsequently processed with a Gaussian blur filter (σ = 0.5). The region of interest (ROI) was defined as the entire region within the tumor, excluding any contrast outside this ROI from analysis. The normalized signal-to-background ratio (SBR) was calculated by dividing the BURST signal within the ROI of the first acquisition by the signal within the same ROI of the second acquisition.

For analyses involving CAR-GV experiments (**Fig. 5** and **Extended Data Figs. 10–11**), BURST images were reconstructed using a correlation-based method (CorrBURST) adapted from recent work^58^, which improves sensitivity and specificity for detecting GV-derived signals *in vivo*. Briefly, CorrBURST identifies pixels exhibiting temporal signal patterns consistent with GV collapse by correlating the observed signal evolution with an expected collapse profile, thereby enhancing GV-specific contrast while suppressing background from non-specific scatterers. A correlation threshold (θ) was applied to identify pixels with GV-specific temporal signatures. We used θ = 0.895 with N = 10 frames, where N denotes the number of frames used for correlation, beginning at the first high-pressure collapse frame. This threshold corresponds to a background rejection rate of approximately 0.9999.

In all images, decibels (dB) was defined as following:

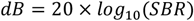

### HLA-I epitope presentation prediction

Prediction of potential HLA class I-presented epitopes was performed using NetMHC4.1 and MUNIS. All possible 9-11 amino acid peptides from each protein sequence were evaluated against the three most frequent human HLA class I alleles, HLA-A*02:01, HLA-A*24:02, and HLA-C*04:01^59^. For MUNIS, peptide predictions were performed using the native protein sequence context. No flanking sequence was provided beyond the termini of each protein.

For MUNIS, peptides with a ligand presentation score greater than 0.8 were classified as predicted presented epitopes. The total number of predicted presented epitopes was then calculated for each protein. For NetMHC4.1, peptide presentation likelihood was evaluated using the eluted ligand rank (EL rank). Peptides with an EL rank below 0.5% for at least one of the evaluated HLA alleles were classified as predicted presented epitopes. The total number of predicted presented epitopes was subsequently calculated for each protein.

### Statistical Analysis

All statistical tests were performed with Prism 10 (Graphpad). All tests were performed with 95% confidence intervals. Standard deviations were checked to make sure appropriate t tests were performed. No other methods were used to determine if data satisfied the assumptions made in the statistical tests.

### Data availability

All plasmids used in this study are available from M.G.S. under a material agreement with the California Institute of Technology. The key genetic constructs will be deposited with Addgene at the time of manuscript publication. Raw data are available from the corresponding author upon reasonable request.

### Code availability

Ultrasound acquisition and processing scripts used to generate key figures and results will be posted to a publicly accessible GitHub repository at the time of manuscript publication.

## ACKNOWLEDGMENTS

The authors thank T. Dilanyan and L. S. Pachter for sharing tissue culture space. Confocal microscopy was performed in the Beckman Institute Biological Imaging Center with help from G. Spigolon. This research was supported by the National Institutes of Health (R01-EB018975 to M.G.S.) and the Chan Zuckerberg Initiative. Related research in the Shapiro laboratory is supported by the Packard Foundation. S.S. was supported by a National Science Foundation Graduate Research Fellowship Program fellowship. M.G.S. is an investigator of the Howard Hughes Medical Institute.

## CONTRIBUTIONS

S.S. conceived and planned the study, generated genetic constructs, and evaluated their performance *in vitro* and *in vivo*. S.S., A.L., and M.S. conducted in vivo experiments. S.S. and M.H.A. designed genetic constructs and established in vitro protocols. J.R. produced lentiviral vectors and performed histology and confocal imaging. B.U.H. conducted in vitro experiments. M.G.S. supervised the study. S.S. and M.G.S. wrote the manuscript with input from A.L., M.H.A., and B.U.H.

## Extended Data Figures

**Extended Data Figure 1:**
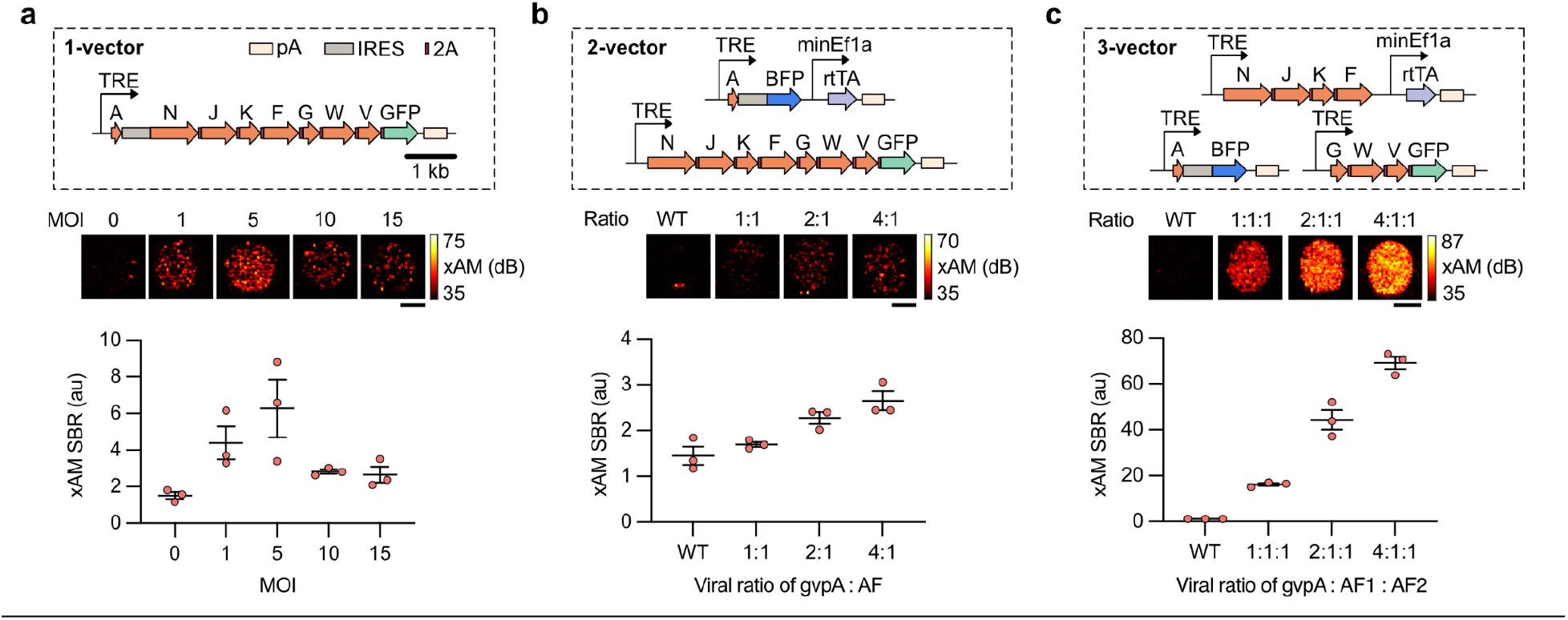
xAM imaging of HEK293T cells expressing GVs using different lentiviral vector designs. **a,** xAM imaging of HEK-TetON cells transduced with the single lentiviral vector encoding all necessary GV genes, shown at increasing MOIs. **b,** xAM imaging of HEK293T cells transduced with the two-vector lentiviral system at increasing ratios of gvpA to assembly factors (AFs). **c,** xAM imaging of HEK293T cells transduced with the three-vector lentiviral system at increasing ratios of gvpA to AF1 and AF2 vectors. Statistical comparisons to non-transduced (WT) controls were performed using Fisher’s LSD test. P-values from left to right: (**a**): 0.0361, 0.0025; (**b**): 0.0065, 0.0007; (**c**): 0.0033, <0.0001, <0.0001. T-values from left to right: (**a**): 2.420, 4.007; (**b**): 3.647, 5.312; (**c**): 4.140, 11.89, 18.73. Df: (**a**): 10; (**b**): 8; (**c**): 8. Significance levels: \**p* < 0.05, \*\**p* < 0.01, \*\*\**p* < 0.001, \*\*\*\**p* < 0.0001; nonsignificant data points are not marked. Error bars represent mean ± s.e.m. of N = 3 biological replicates. Each data point reflects the arithmetic mean of N = 2 technical replicates. Scale bars for all ultrasound images represent 1 mm. au: arbitrary units.

**Extended Data Figure 2:**
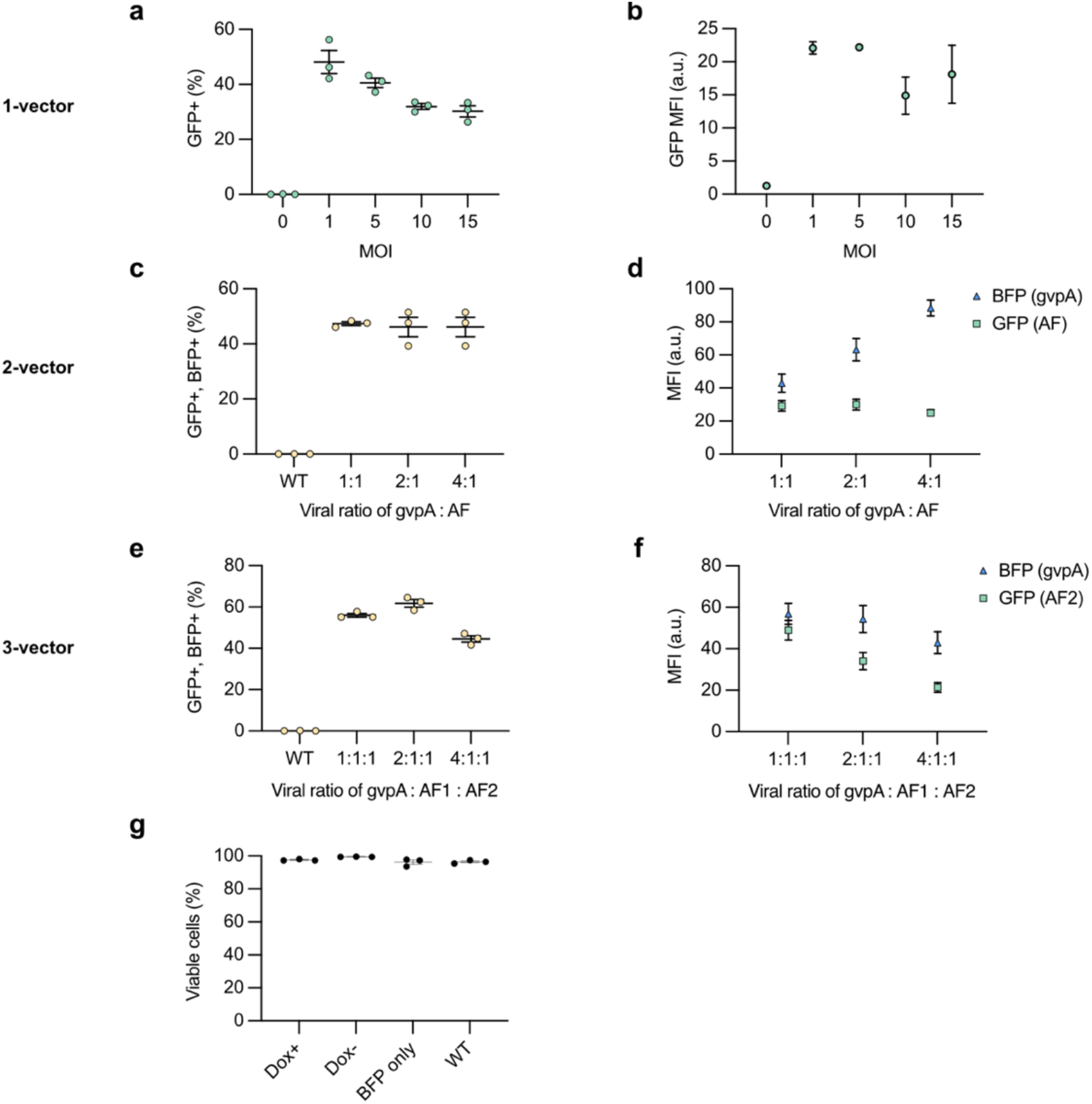
Flow cytometry characterization of HEK293T cells transduced with various lentiviral designs encoding GV genes. **a-b**, Percentage of HEK-TetON cells transduced with the single-vector virus that express GFP, the marker for transduction and their mean fluorescence intensity (MFI). **c-d**, Percentage of HEK293T cells transduced with the two-vector lentivirus system, expressing both GFP and BFP, markers for double transduced cells, and their MFI. **e-f**, Percentage of HEK293T cells transduced with the three-vector lentivirus system expressing GFP and BFP, markers for triple transduced cells, and their MFI. **g**, Percentage of live cells transduced with the three-vector lentivirus system at a 4:1:1 ratio (gvpA : AF1 : AF2) and sorted for triple-transduction, following 3 days of Dox induction, compared to cells transduced with a control virus constitutively expressing BFP at the same total MOI of 10, and WT cells. Statistical comparisons were made using Fisher’s LSD test comparing the GV-expressing cells (Dox+) to uninduced cells (Dox-), the BFP control line, and wild-type (WT) cells. All p-values were non-significant. P-values from left to right: 0.1017, 0.2189, 0.3020. Error bars represent mean ± s.e.m. of N = 3 biological replicates.

**Extended Data Figure 3:**
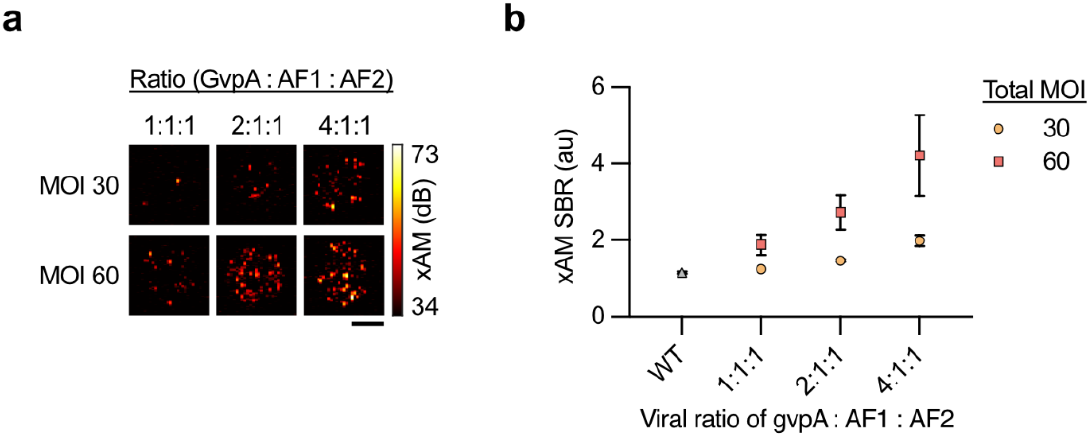
xAM imaging of Jurkat cells transduced with the 3-vector lentivirus system at varying MOIs. **a-b**, xAM imaging of Jurkat cells transduced with three lentiviral vectors encoding all necessary GV genes, shown at different vector ratios and total MOIs of 30 and 60. Statistical comparisons for panel (**b**) were performed using Fisher’s LSD test comparing each condition to WT. P-values smaller than 0.5: MOI 60 (2:1:1): 0.0257; MOI 60 (4:1:1): 0.0003. Error bars represent mean ± s.e.m. N = 3 biological replicates. Each data point represents the arithmetic mean of N = 2 technical imaging replicates. All ultrasound image scale bars represent 1 mm.

**Extended Data Figure 4:**
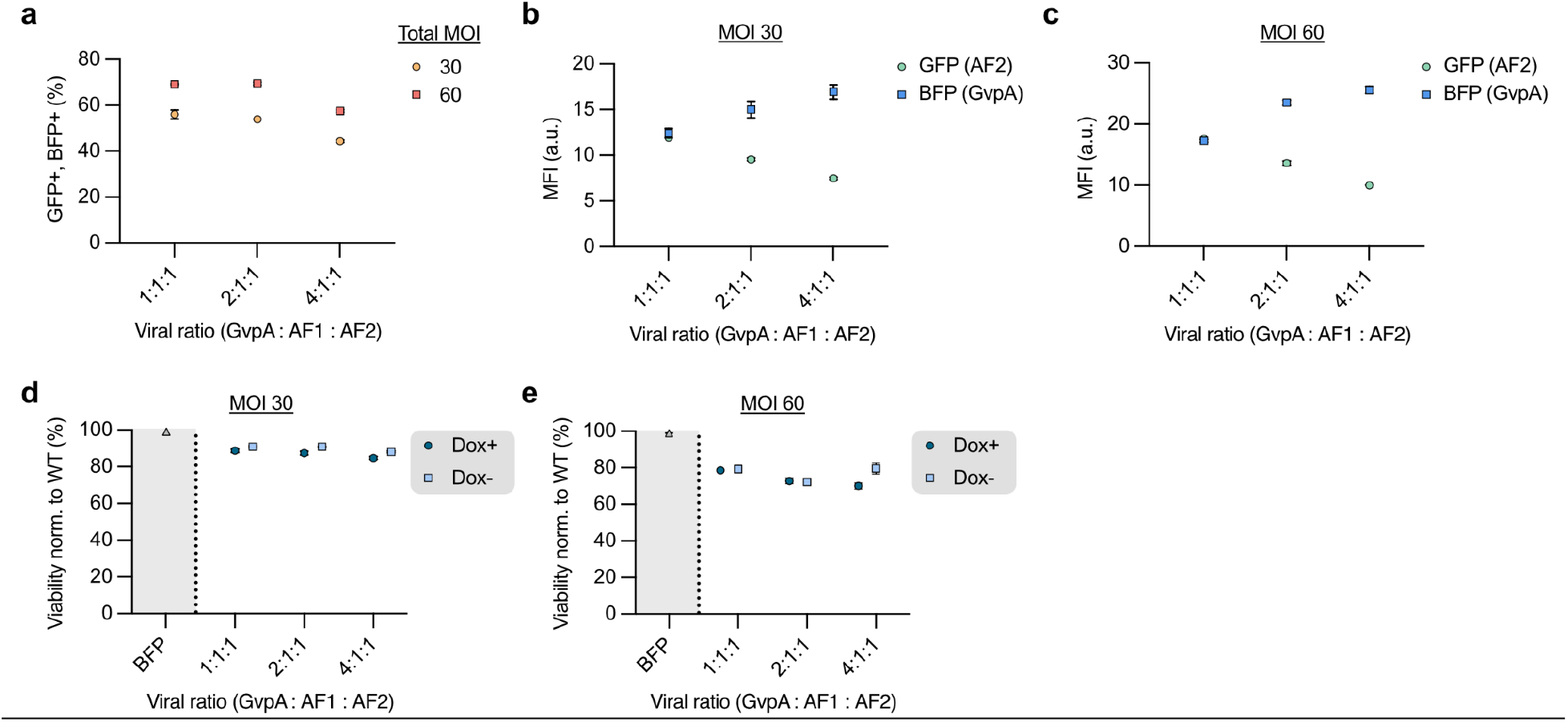
Flow cytometry analysis of Jurkat cells expressing GVs. **a**, Percentage of Jurkat cells transduced with all three GV lentiviruses that express both GFP and BFP. **b-c**, Mean fluorescence intensity (MFI) of GFP and BFP expression in the cells from panel (**a**). **d-e**, Cell viability of Jurkat cells transduced with GV-expressing lentiviruses at MOIs of 30 (**d**) and 60 (**e**), with and without Dox induction, and Jurkat cells transduced with a virus constitutively expressing BFP at the same MOIs. Error bars represent mean ± s.e.m. of N = 3 biological replicates.

**Extended Data Figure 5:**
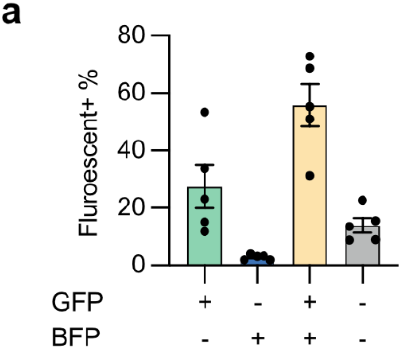
Characterization of sorted T cell subpopulations using fluorescent transduction markers. **a**, Flow cytometry analysis of Dox-induced and sorted T cells shows that the majority of T cells express both lentiviral transduction markers, GFP and BFP, four days after sorting. As the cells were sorted mainly based on GFP expression (assembly factor viral vectors), a smaller subpopulation expresses only GFP without BFP, the marker for the gvpA-encoding viral vector. Error bars represent mean ± s.e.m. of N = 5 PBMC donors.

**Extended Data Figure 6:**
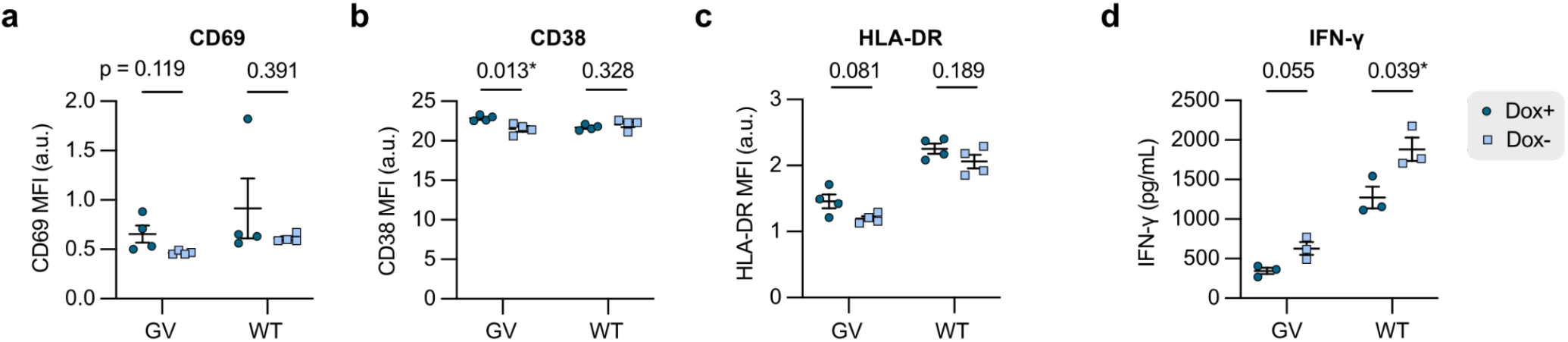
GV expression does not affect T cell activity markers. MFI of **a,** CD69, **b,** CD38 and **c,** HLA-DR in T cells transduced with GV-encoding 3-vector lentiviral system and in untransduced WT T cells, with and without Dox induction (Dox+ and Dox-). N = 4 replicates. **d,** IFN-γ concentration measured in T cell culture supernatants. N = 3 replicates. Two-tailed Welch’s t test. P-values are stated above data points. Error bars represent mean ± s.e.m.

**Extended Data Figure 7:**
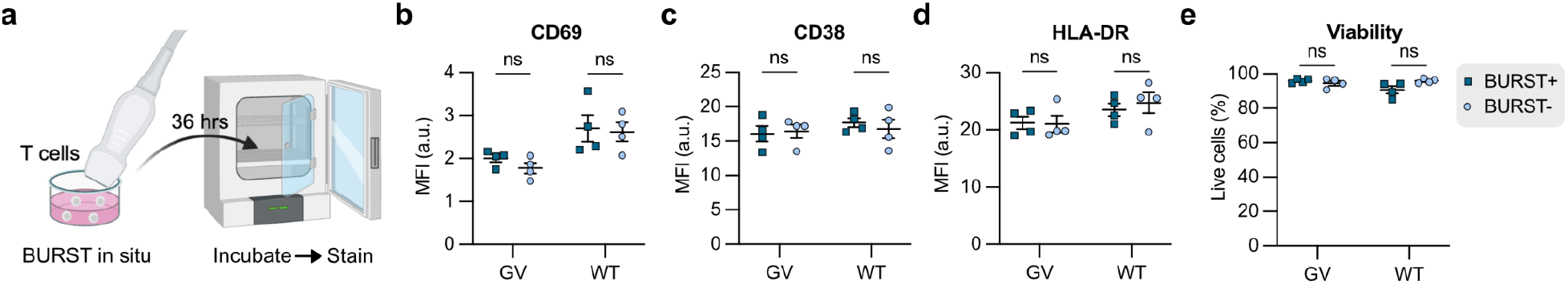
BURST imaging does not affect T cell activity state or viability. **a,** Schematic of the experiment in which Dox-induced GV expressing and WT T cells in suspension culture were exposed to BURST imaging and subsequently incubated at 37°C for additional 36 hours before staining for activity and viability markers. MFI of **b,** CD69, **c,** CD38, and **d,** HLA-DR. e, Percentage of viable cells (7AAD-). N = 4 replicates. Two-tailed Welch’s t test. ns: p ≥ 0.05. Error bars represent mean ± s.e.m.

**Extended Data Figure 8:**
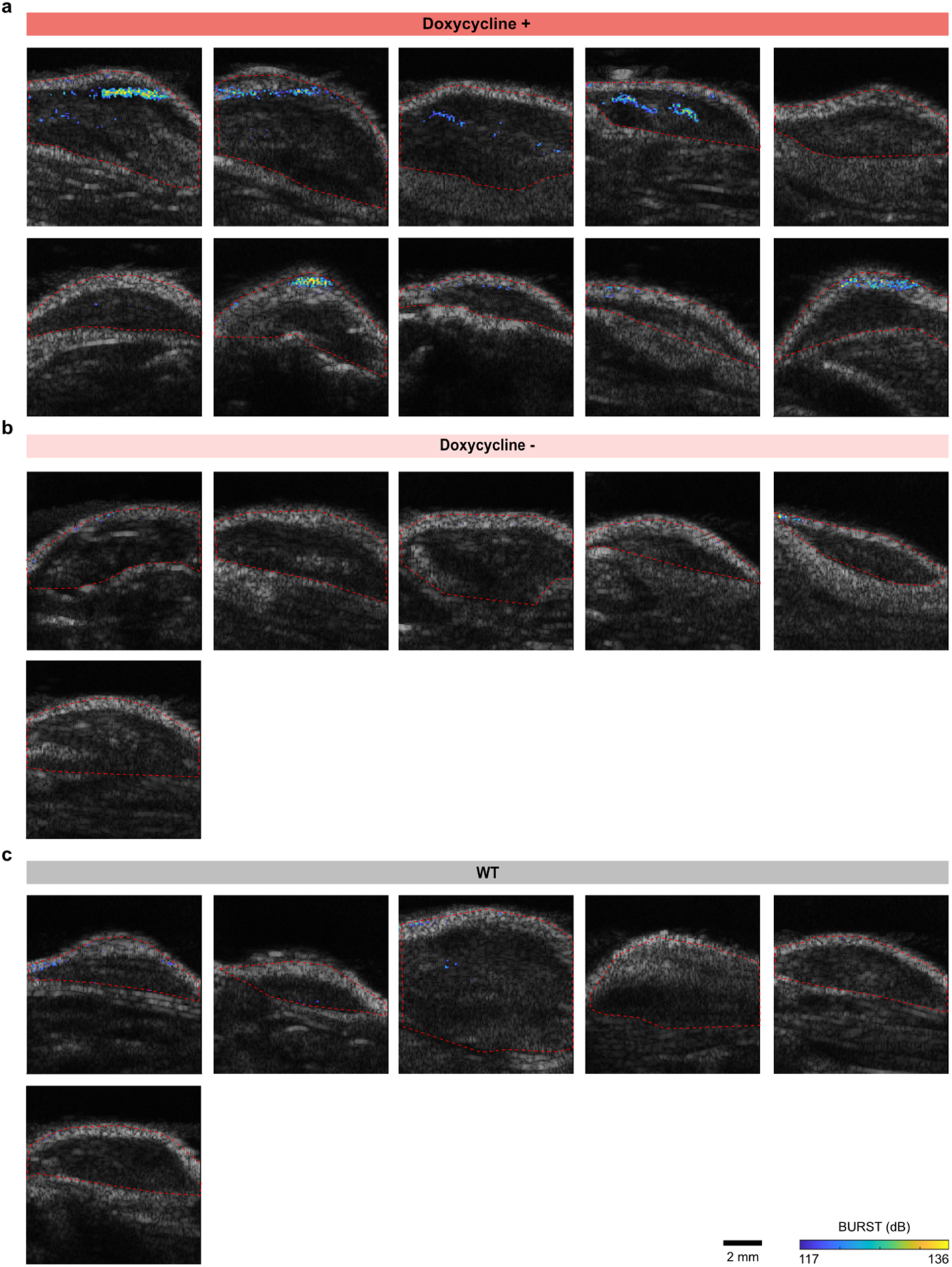
Ultrasound imaging of Raji tumors infiltrated with T cells. Representative BURST images (colormap) overlaid on anatomical B-mode images (grayscale) of subcutaneous Raji tumors, with each image representing a different mouse. **a**, Tumors from mice infused with virally transduced T cells, induced with doxycycline for 3 days to express GVs. **b**, Tumors from mice infused with transduced T cells but left uninduced. **c**, Tumors from mice infused with wild-type (WT) T cells.

**Extended Data Figure 9:**
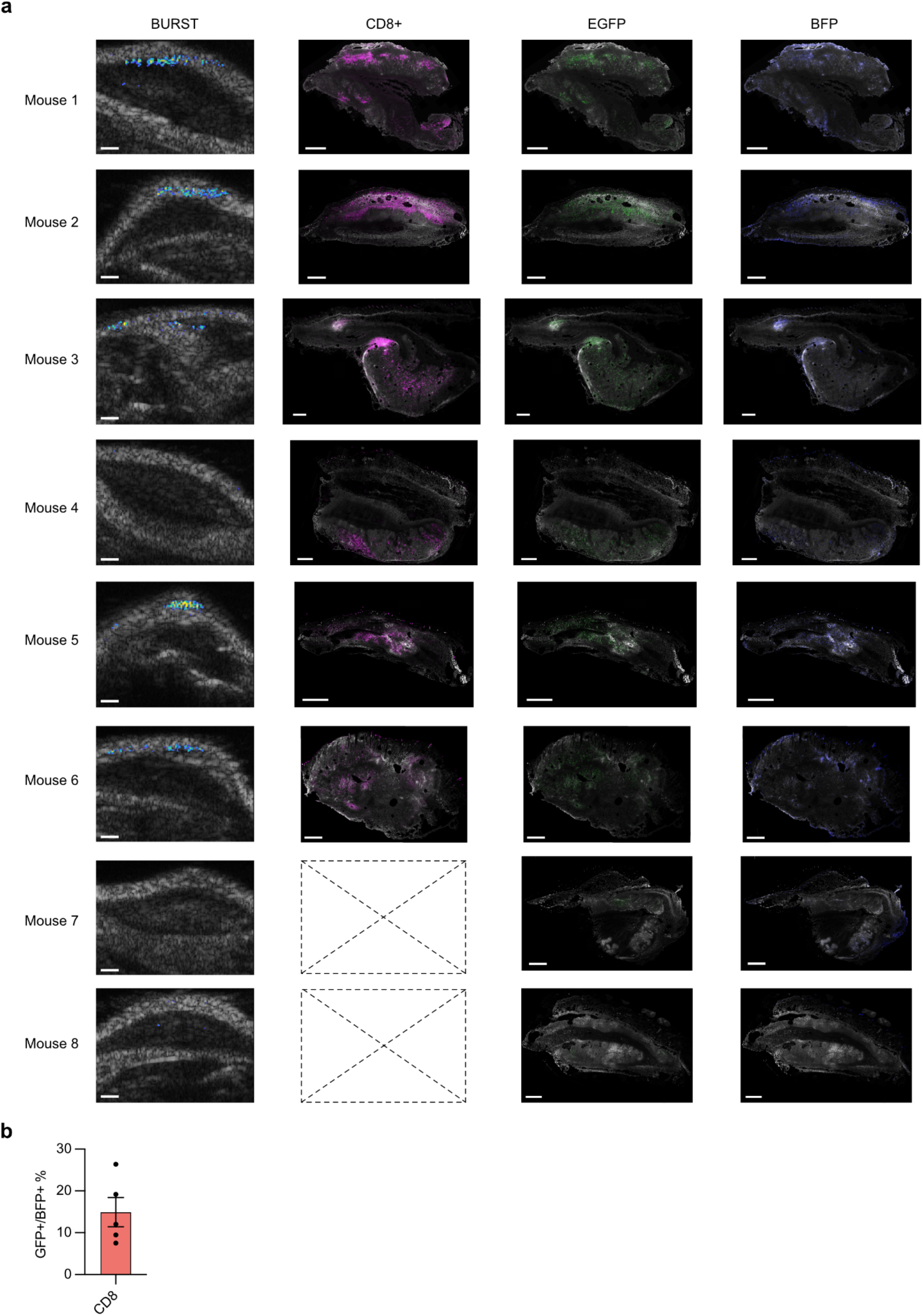
Assessing T cell infiltration in tumors. **a**, Ultrasound images (BURST overlaid on B-mode) of Dox-induced, GV-expressing T cells in Raji-Antares tumors, alongside corresponding histology showing cytotoxic CD8+ T cells, GFP+ T cells, and BFP+ T cells. Mice 7 and 8 were not processed for anti-CD8 staining due to a negligible presence of GFP/BFP-expressing T cells. Note that the histology and ultrasound images are not an exact spatial match, but are within 1 mm of each other. White: Antares; Magenta: anti-CD8; Green: GFP; Blue: BFP. All scale bars represent 1 mm. **b,** Percentage of CD8+ T cells that were GFP+/BFP+ (triple-transduced population) as determined by flow cytometry of digested tumors. Error bars represent mean ± s.e.m.

**Extended Data Figure 10:**
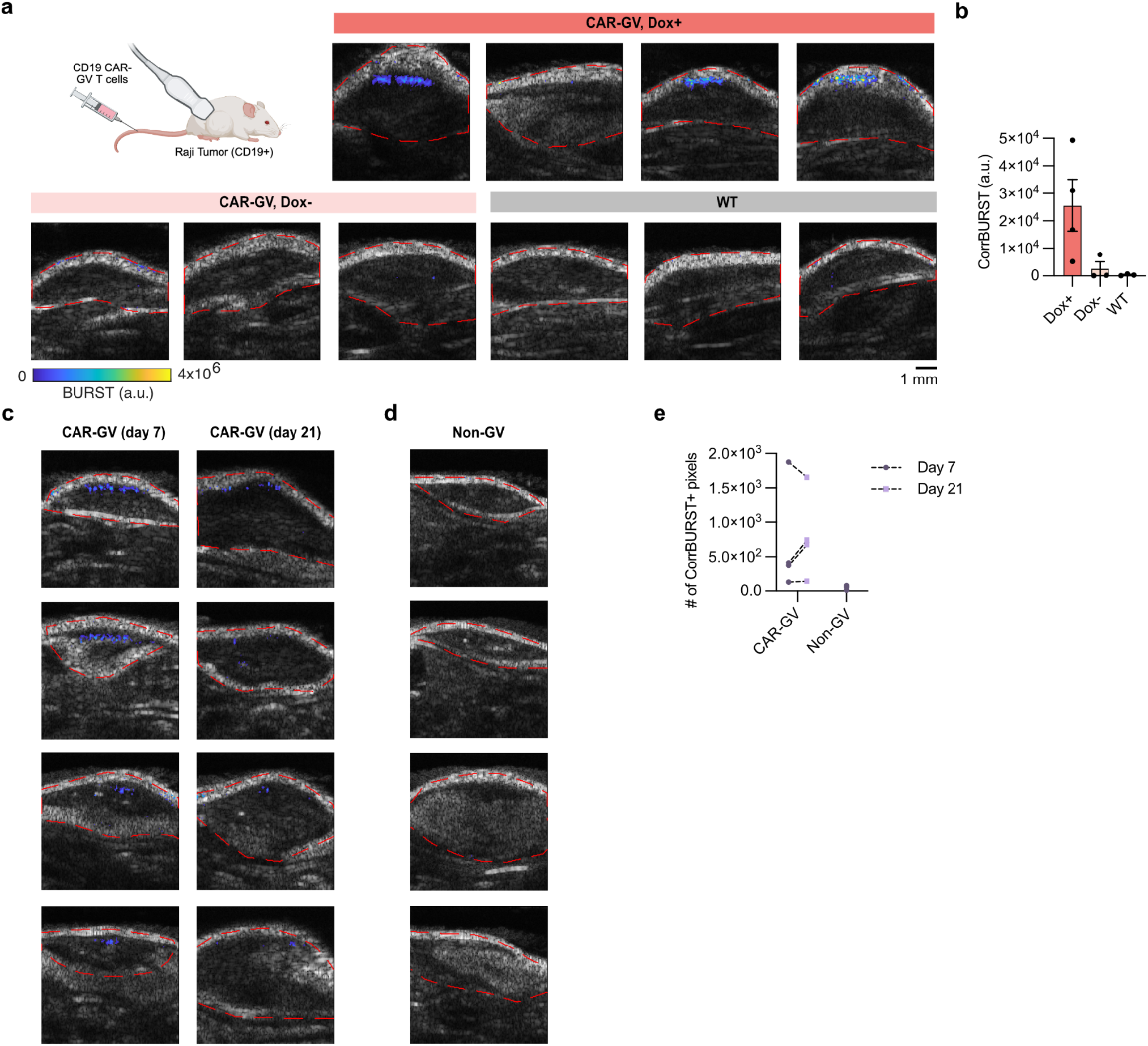
Longitudinal imaging of CAR-GV T cell homing to tumors. **a,** Representative BURST images of tumors from NSG mice bearing subcutaneous Raji–Antares tumors following systemic injection of CAR-GV–transduced T cells or untransduced WT T cells; images were acquired 7 days after T cell injection, red lines indicate tumor boundaries used for signal quantification, and each image corresponds to a different mouse. **b,** CorrBURST signal intensity quantified within tumors shown in (a). **c,** NSG mice were systemically injected with CAR-GV T cells or **d,** non-GV–expressing T cells and imaged longitudinally on days 7 and 21 after T cell injection; images on the same row in (c) are from the same mouse; representative images from each mouse are shown. **d,** CorrBURST signal quantified over the entire tumor volume for each mouse, plotted as the sum of CorrBURST-positive pixels per tumor.

**Extended Data Figure 11:**
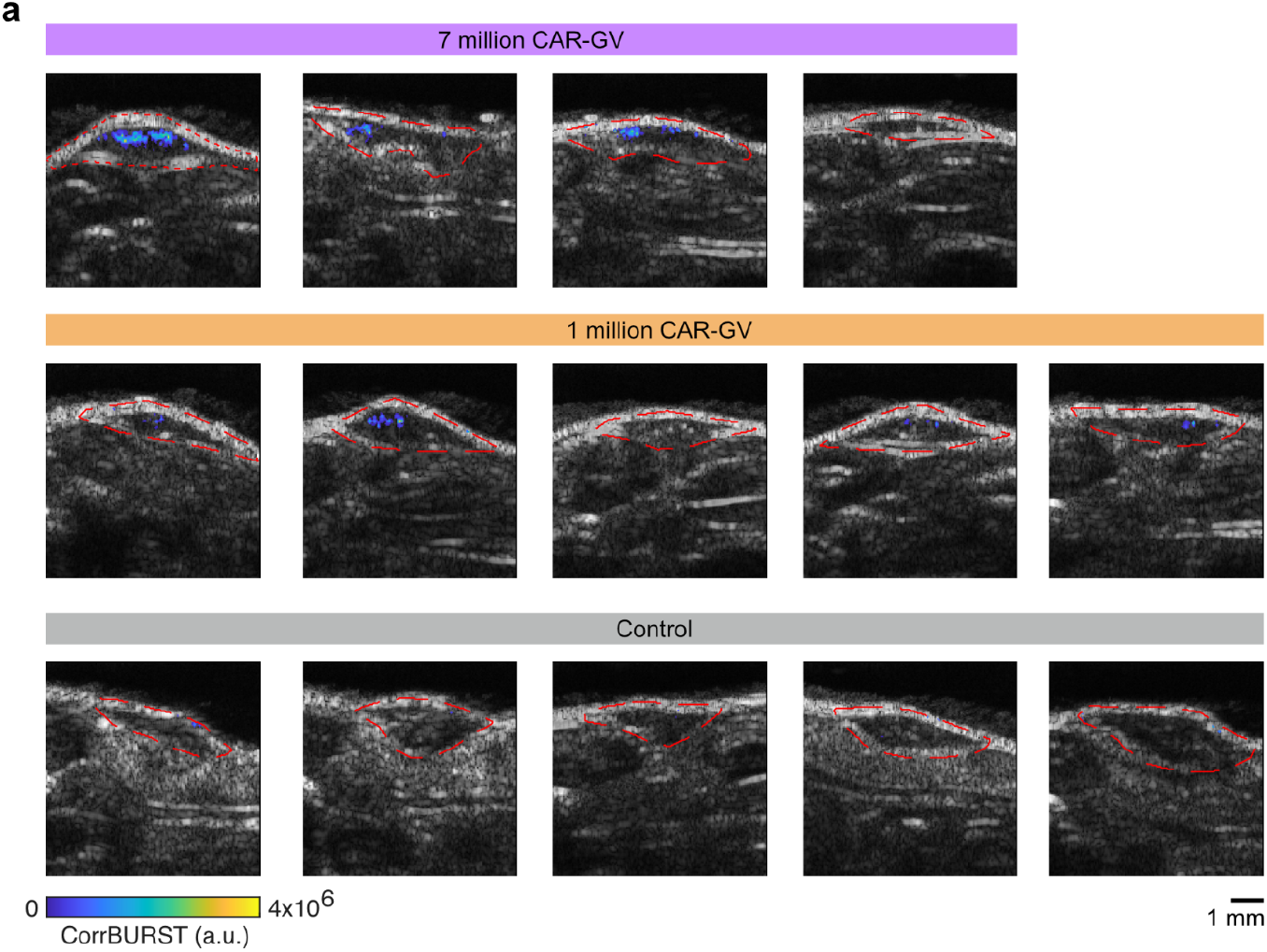
BURST imaging of CAR-GV T cells in tumors following systemic injections of different T cell doses. **a,** Representative day 11 CorrBURST images (colorbar) overlaid on anatomical B-mode images (grayscale) of Raji tumors in mice injected with Dose A (7 × 10⁶) or Dose B (1 × 10⁶) CAR-GV T cells, or with non-GV–expressing T cells (control); red lines indicate tumor boundaries used for signal quantification. Each image corresponds to a different mouse, and the plane with the highest CorrBURST signal from each mouse is shown.

**Extended Data Figure 12:**
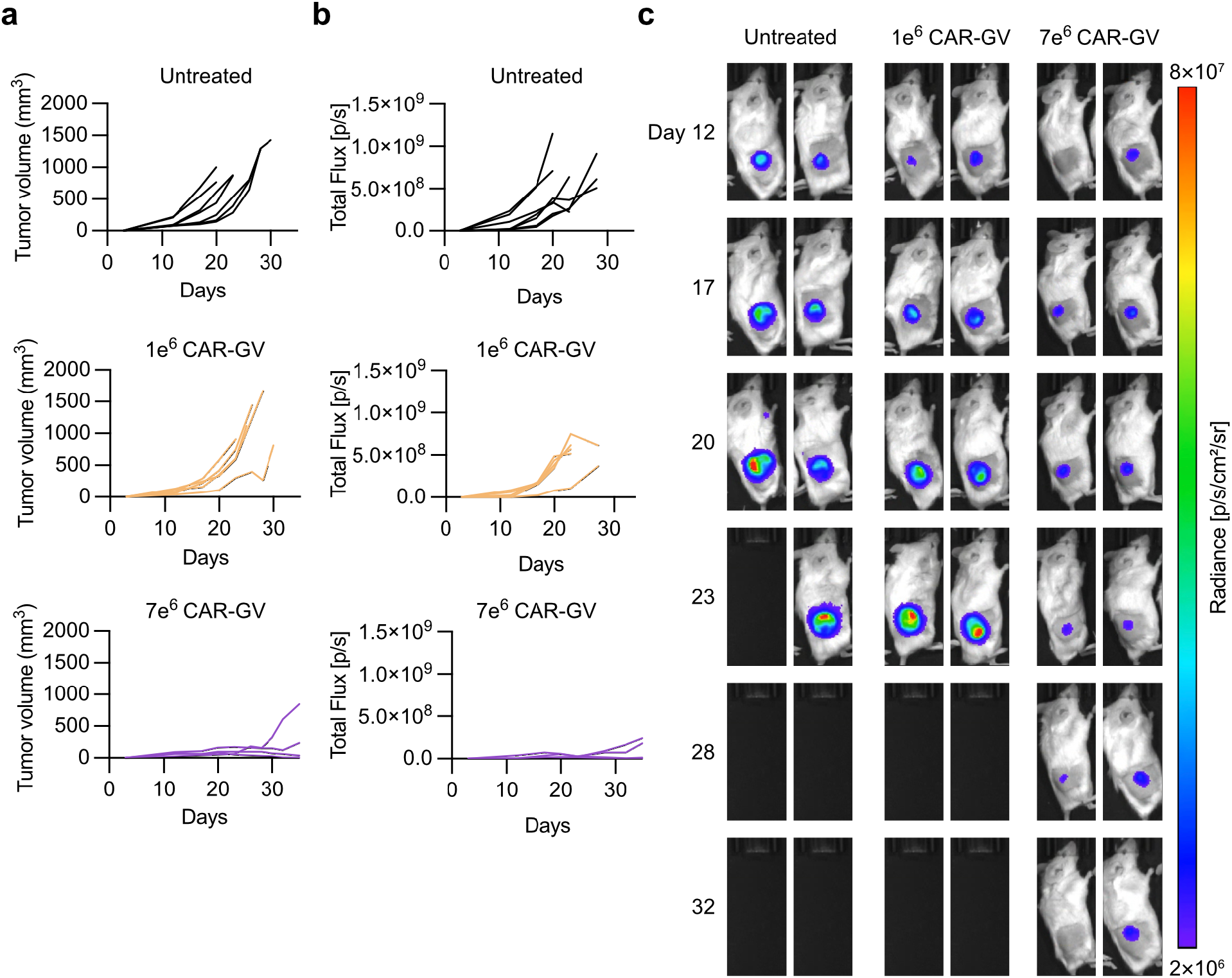
Tumor size measurements. **a,** Tumor volumes for individual mice over time, measured with a caliper. Each line represents a single mouse. **b,** Total flux from tumors for individual mice over time, measured by IVIS. Each line represents a single mouse. **c,** Representative IVIS images of N = 2 mice from each group.

**Extended Data Figure 13:**
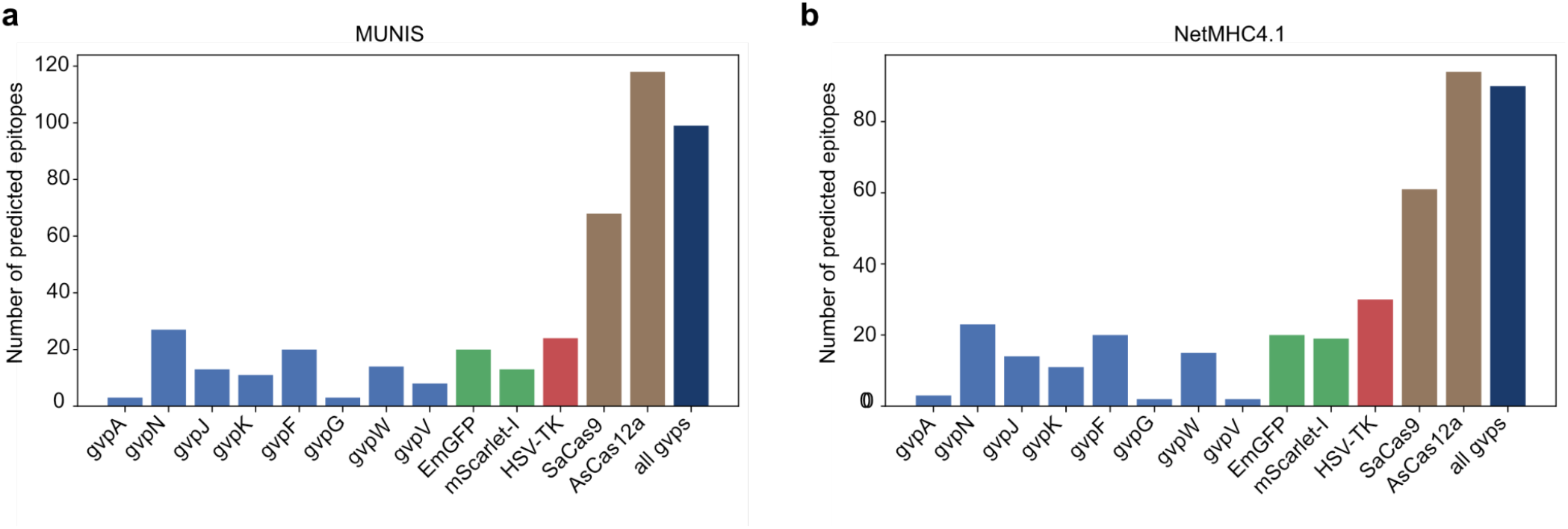
In silico prediction of HLA class I presentation burden across GV proteins, established reporter genes, and Cas proteins. **a,** Total number of MUNIS epitope predictions per protein. **b,** Total number of NetMHC epitope predictions per protein.

## Supplementary Figures

**Supplementary Figure 1:**
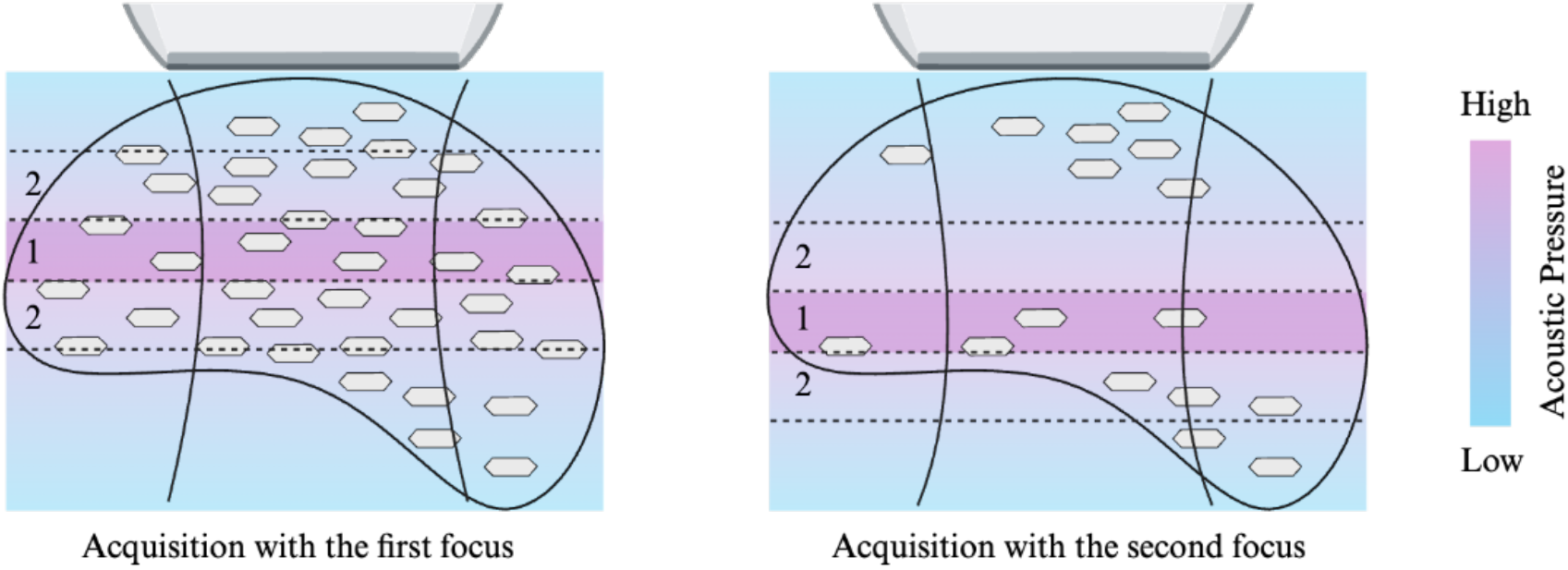
Out-of-focus partial collapse in BURST imaging with a linear array transducer. BURST imaging with a linear array transducer generates a focused ultrasound beam, with the highest acoustic pressure at the center of the focus (zone 1). One limitation of this method is the potential for collapsing GVs outside the focus of imaging (zone 2) at lower pressures, without producing a detectable BURST signal. While varying the focus to different depths allows us to capture some of the collapse-based nonlinear scattering, this method of imaging may not be perfectly capturing GV expression throughout the entire tumor. A potential solution would be to use a matrix array or row-column array transducer, which can generate a consistent axial pressure field along the ultrasound beam.

**Supplementary Figure 2:**
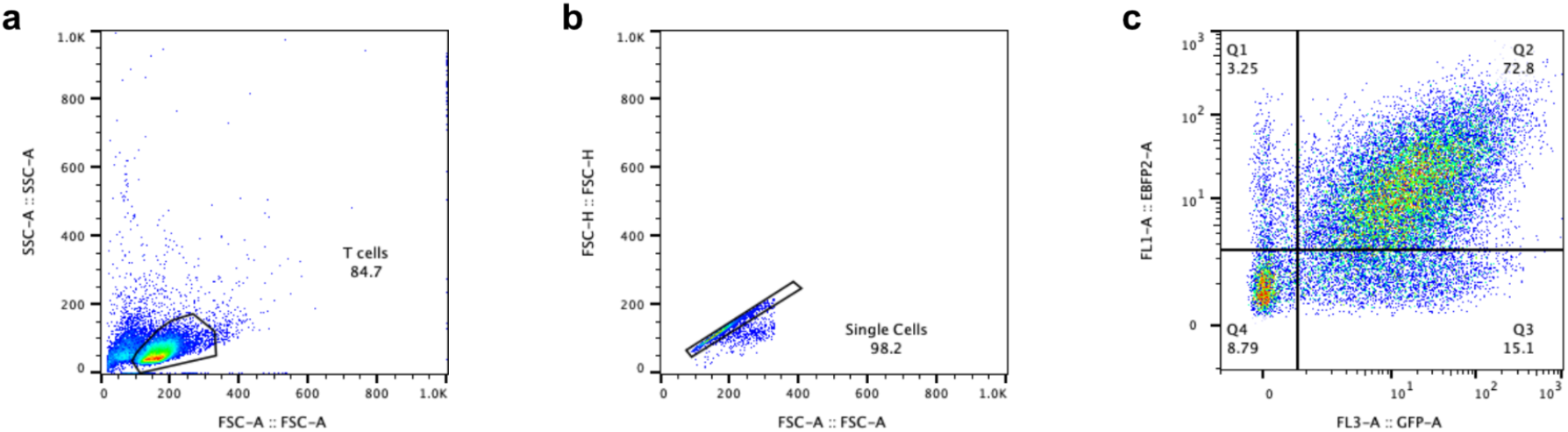
Flow cytometry gating strategy. Flow cytometry analysis of representative T cells transduced with the 3-vector GV expressing lentiviral vectors and induced with doxycycline. **a**, Cells are selected based on their FSC-SSC scattering. **b**, Within the “T cells” subpopulation, single cells are selected based on their FSC-H and FSC-A scattering. **c**, Within the “Single Cells” subpopulation, a threshold for each fluorescence channel is chosen based on wild-type/unstained controls. The thresholds are applied to all samples within a single experiment.

---

**Supplementary Table 1.**
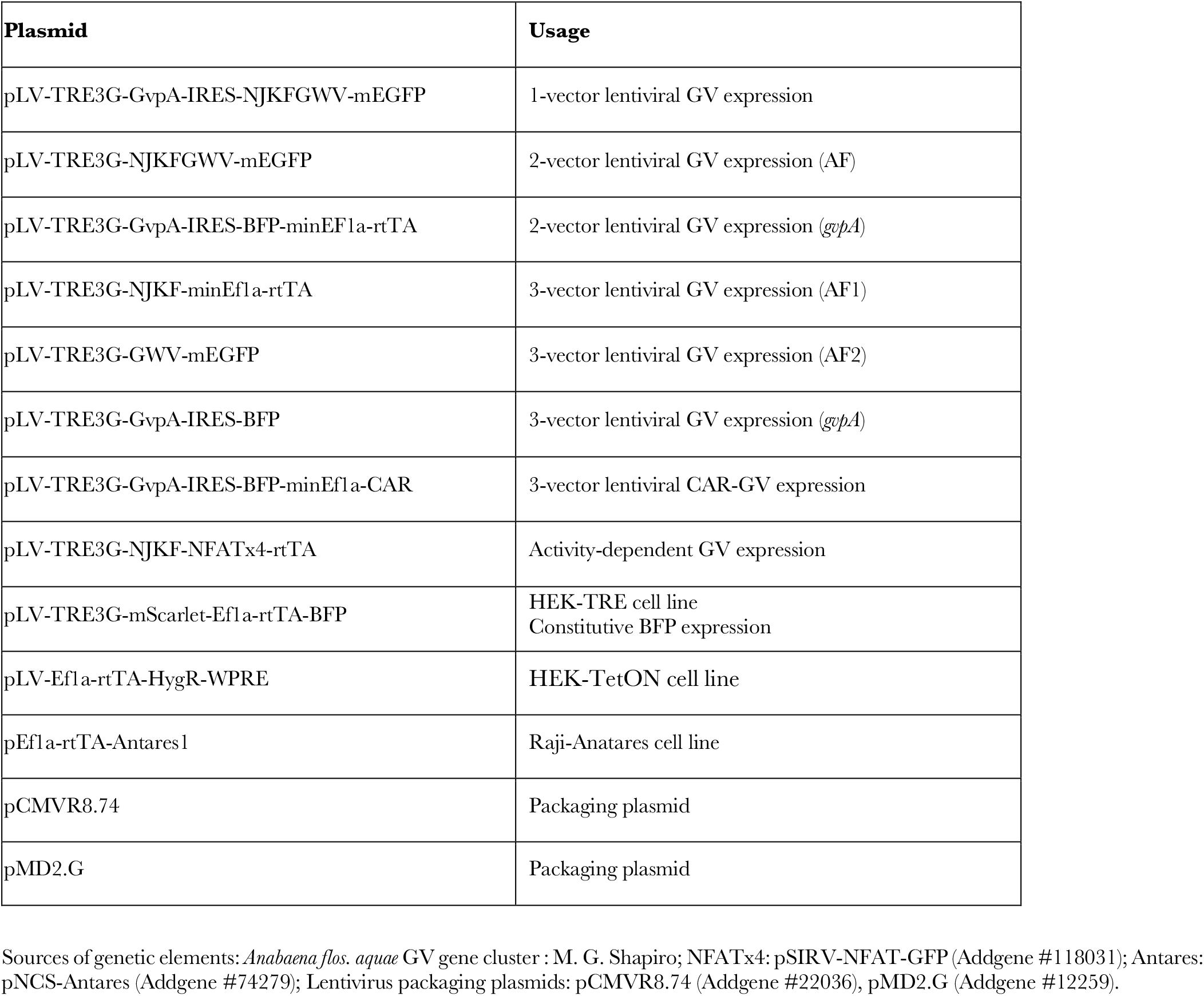
Genetic constructs used in the study.

**Supplementary Table 2.**
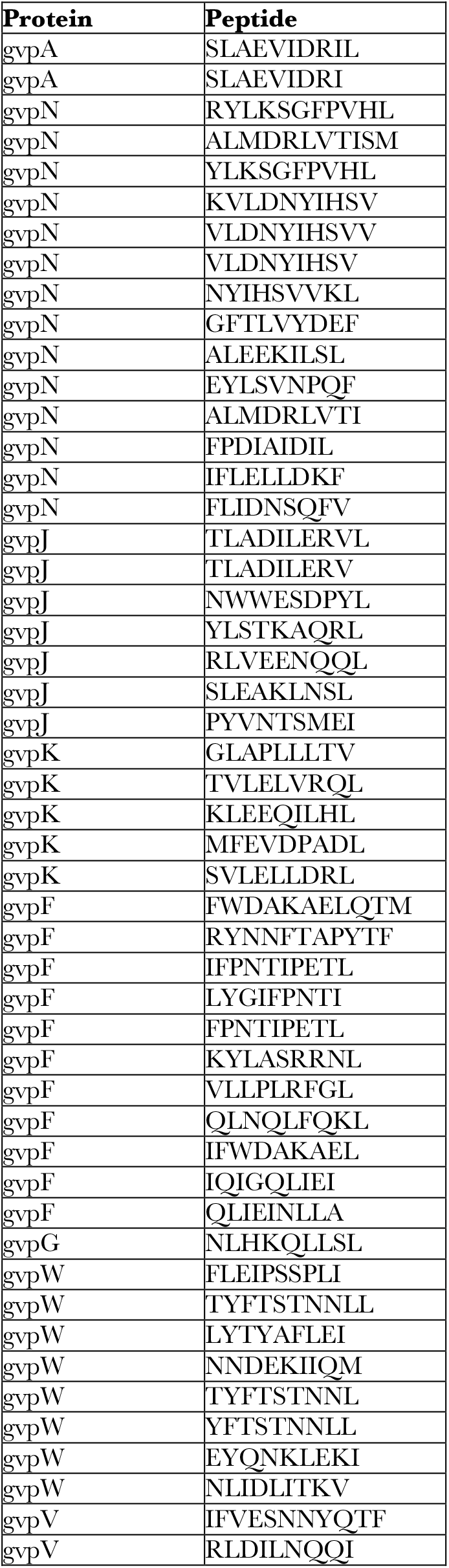
Predicted HLA class I-presented peptides from GV proteins identified by both MUNIS and NetMHCpan 4.1.

